# Electron cryo-microscopy of Bacteriophage PR772 reveals the composition and structure of the elusive vertex complex and the capsid architecture

**DOI:** 10.1101/645523

**Authors:** Hemanth. K. N. Reddy, Janos Hajdu, Marta Carroni, Martin Svenda

## Abstract

Bacteriophage PR772, a member of the *Tectiviridae* family, has a 70-nm diameter icosahedral protein capsid that encapsulates a lipid membrane, dsDNA, and various internal proteins. An icosahedrally averaged CryoEM reconstruction of the wild-type virion and a localized reconstruction of the vertex region reveals the composition and the structure of the vertex complex along with new protein conformations that play a vital role in maintaining the capsid architecture of the virion. The overall resolution of the virion is 2.75 Å, while the resolution of the protein capsid is 2.3 Å. The conventional penta-symmetron formed by the capsomeres is replaced by a large vertex complex in the pseudo T=25 capsid. All the vertices contain the host-recognition protein, P5; two of these vertices show the presence of the receptor-binding protein, P2. The 3D structure of the vertex complex shows interactions with the viral membrane, indicating a possible mechanism for viral infection.

## Introduction

Bacteriophage PR772 is a double-stranded DNA (dsDNA) virus with a 70 nm icosahedral protein capsid encapsulating the internal lipid bilayer along with numerous proteins and a 15 kbp long linear genome. It belongs to the *Tectiviridae* family and infects gram negative hosts like *Escherichia coli*, *Salmonella typhimurium* and other bacteria, carrying a R772 plasmid-encoded receptor complex through which DNA can be transported during bacterial conjugation (J. N. Coetzee, Lecatsas, Coetzee, & Hedges, 1979; W. F. Coetzee & Bekker, 1979; Lute, Aranha, & Tremblay, 2004; H. K. N. Reddy et al., 2017). Much of the functional knowledge about the viral proteins is inferred from previous studies on PRD1, a close relative of PR772 (Lute et al., 2004). The most striking features of phages from those members of the *Tectiviridae* family that infect Gram-negative bacteria, is the presence of an inner lipid membrane and lack of a tail in the dormant viral particle. During the process of infection, these viruses produce a membranous tube derived from the inner membrane of the viral particle, lined from the inside by proteins P7, P11 and P32 (Peralta et al., 2013; Saren et al., 2005). This tube is used to inject the viral dsDNA into the host (Peralta et al., 2013; Saren et al., 2005).

Genome analysis of PR772 identified 32 open reading frames (ORFs) containing at least 40 codons (Lute et al., 2004). Twenty-eight annotated proteins are known to be expressed from the genome of which 3 do not make it into the final assembly (Butcher, Manole, & Karhu, 2012; Lute et al., 2004). In previous studies on PRD1, a close relative of PR772, it was shown that the capsid is formed by proteins P3 (major capsid protein), P30 (capsid associated protein) and the vertex complex. The vertex complex, includes the penton protein (P31), the host-recognition protein (P5) and receptor-binding proteins (P2) along with the infectivity protein (P16) that acts as a cementing protein which holds the vertex complex together and also stabilizes the pseudo-icosahedral capsid of the virus. Proteins P6, P9, P20 and P22 are involved in DNA packaging and proteins P7, P14, P11, P18, P32 and P34 are responsible for DNA delivery to the host (A Marika Grahn, Daugelavicius, & Bamford, 2002; Lute et al., 2004). The *Tectiviridae* family of viruses are also known to have structural similarities to adenovirus (Saren et al., 2005).

In previous studies on bacteriophage PRD1, a very close relative to PR772 from the *Tectiviridae* family, many models have been proposed to explain the architecture of the penton base and the vertex complex (Huiskonen, Manole, & Butcher, 2007; Javier Caldentey, Roman Tuma, & Bamford, 2000; Rydman et al., 1999; Sokolova et al., 2001). With the available experimental data, the generally accepted composition of the vertex complex of *Tectiviridae* is that P31 is a homopentamer and forms the penton base. P5 and P2 are attached to the P31 penton base but there arrangement is not known. (Butcher et al., 2012; Huiskonen et al., 2007). The high resolution x-ray crystallographic structure of the P2 protein is known but the location and orientation with respect to the penton base in its functional form is currently not known (Xu, Benson, Butcher, Bamford, & Burnett, 2003). Putatively, it is suggested that the beta-propeller motif of P2 might be involved in binding to the host receptor.

Here we present the high-resolution structure of bacteriophage PR772 using electron cryo-microscopy (CryoEM). The high-resolution map helped us to resolve subtle variations in the protein conformations and their influence on the formation of a viable viral particle. The N-terminal region of the P3 subunits have 3 conformations, not two as previously believed (Abrescia et al., 2004). The new N-terminal P3 conformation plays an important role in accommodating the P30 protein during particle assembly. The C-terminal region of P3 not only shows the formation of a β-sheet with P30 but also helps in locking the adjacent trisymmetron through a hinge mechanism, thus facilitating the formation of icosahedral particles and regulating their size. Localized asymmetric reconstruction of the vertex region of PR772 revealed a P5-P31 heteropentameric base and the binding of P2 to P5 in the complex. A combination of high-resolution icosahedral symmetrized single-particle reconstruction and localized asymmetric reconstruction has enabled us to answer some of the intriguing questions about the particle architecture, composition of the penton base and arrangement of the vertex complex.

## Results

### High resolution capsid map at 2.3 Å

The structure of bacteriophage PR772 was determined by electron cryo-microscopy. The overall resolution determined by Fourier shell correlation (FSC) @0.143 was 2.75 Å (Figure 1–figure supplement 1). The local resolution estimated using the two unfiltered final half maps with ResMap (Kucukelbir, Sigworth, & Tagare, 2014) showed that most of the capsid was resolved to 2.30 Å (Figure 1A and B). The resolution of the regions that interacts with the inner lipid bilayer was lower at about 3.2 Å (Figure 1C). The areas around the icosahedral five-fold axes had resolutions varying between 2.3 – 3.0 Å (Figure 1 D-F). At a root mean square (RMS; deviation away from noise as visualised in Coot, where noise is 0) of 4.2, the side chains of the amino acid residues were visible for most of the capsid region. The inner membrane layers of lipid, protein and dsDNA were smeared due to averaging and symmetry mismatch. The resolution in most of the regions was high enough (3.2 Å – 2.3 Å) to build a de-novo model of the asymmetric unit comprising of P3 (Figure 2–figure supplement 1), P30 (Figure 2–figure supplement 2), P5 (Figure 4–figure supplement 3), P31 (Figure 4–figure supplement 3) and P16 (Figure 6–figure supplement 1) into the icosahedrally averaged CryoEM map (Figure 1H and Table 1).

**Figure 1.**
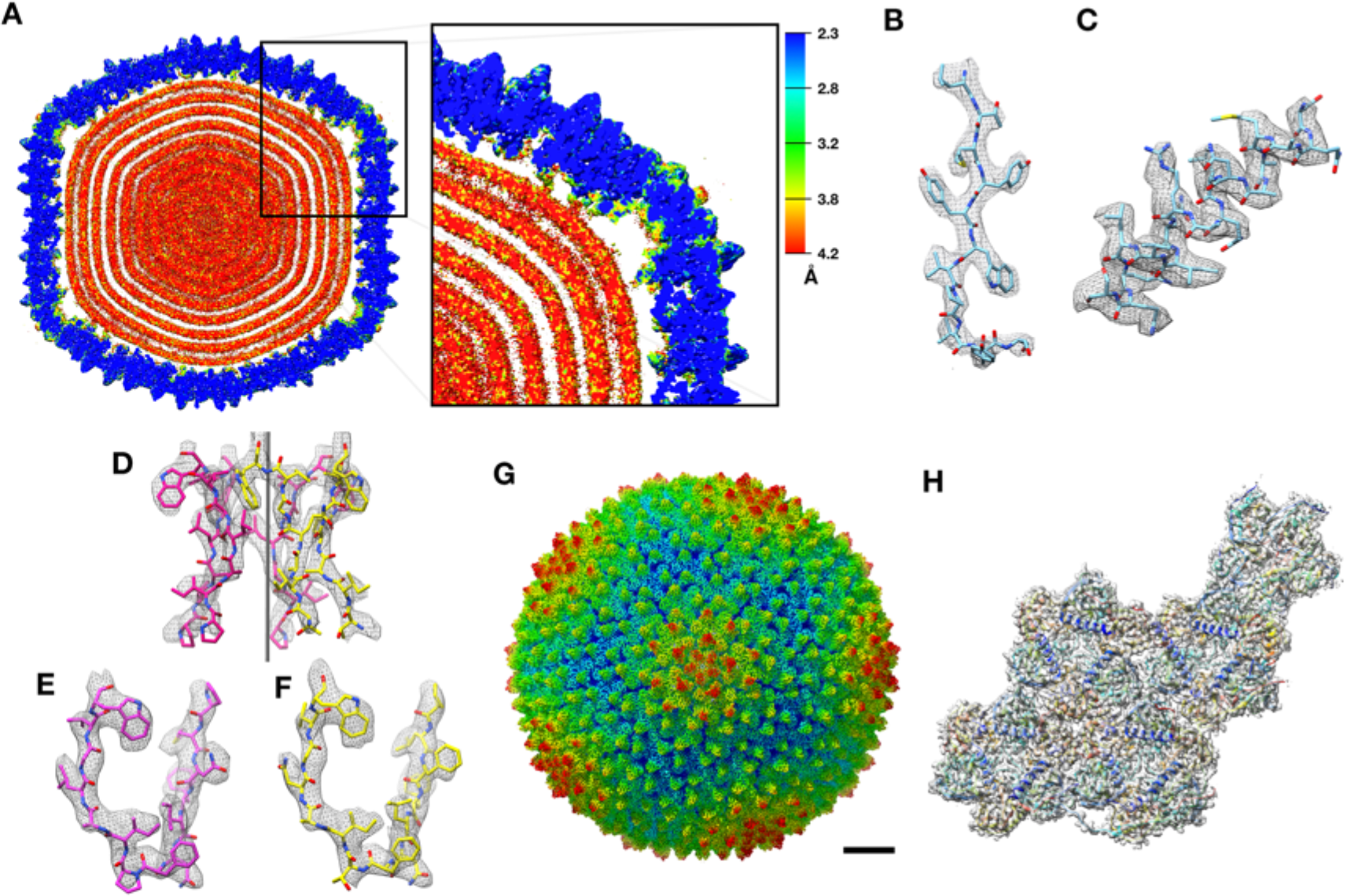
CryoEM structure of PR772 and resolution estimates. (**A**) The local resolution estimate of the CryoEM map from ResMap. The map shows the distribution of resolution in different regions. (Visualized using USCF Chimera, with volume viewer parameters: Style surface, step 1 and level 0.037, Plane 418, Axis Y, Depth 23). Most of the capsid that was used for model building is resolved at 2.3 Å. (**B, C and D**) Show the quality of the map at different regions. (**B**) Quality of the map at the core of the capsid protein P3 (Chain B, residues 162-173) where the local resolution estimate is 2.3 Å. **(C**) Quality of the map close to the membrane (P3 Chain B, residues 18-35) where the resolution is estimated at 3.2 Å. **(D**) Quality of the map close to the five-fold vertex of the icosahedral viral particle. The black vertical line represents the five-fold axis. (**E and F**) The initial model fit of P5 residues 108-121(in pink) and P31 residues 113-126 (in yellow) to the same region of the map using Phenix: Find helix and sheets with respective protein sequences as input. (**G**) The post processed map of PR772 and the scale bar represents 10 nm. (**H**) The map:model fit of the asymmetric unit as seen from the inside of the viral particle. *Also refer* Figure 1–figure supplement 1

**Figure 2:**
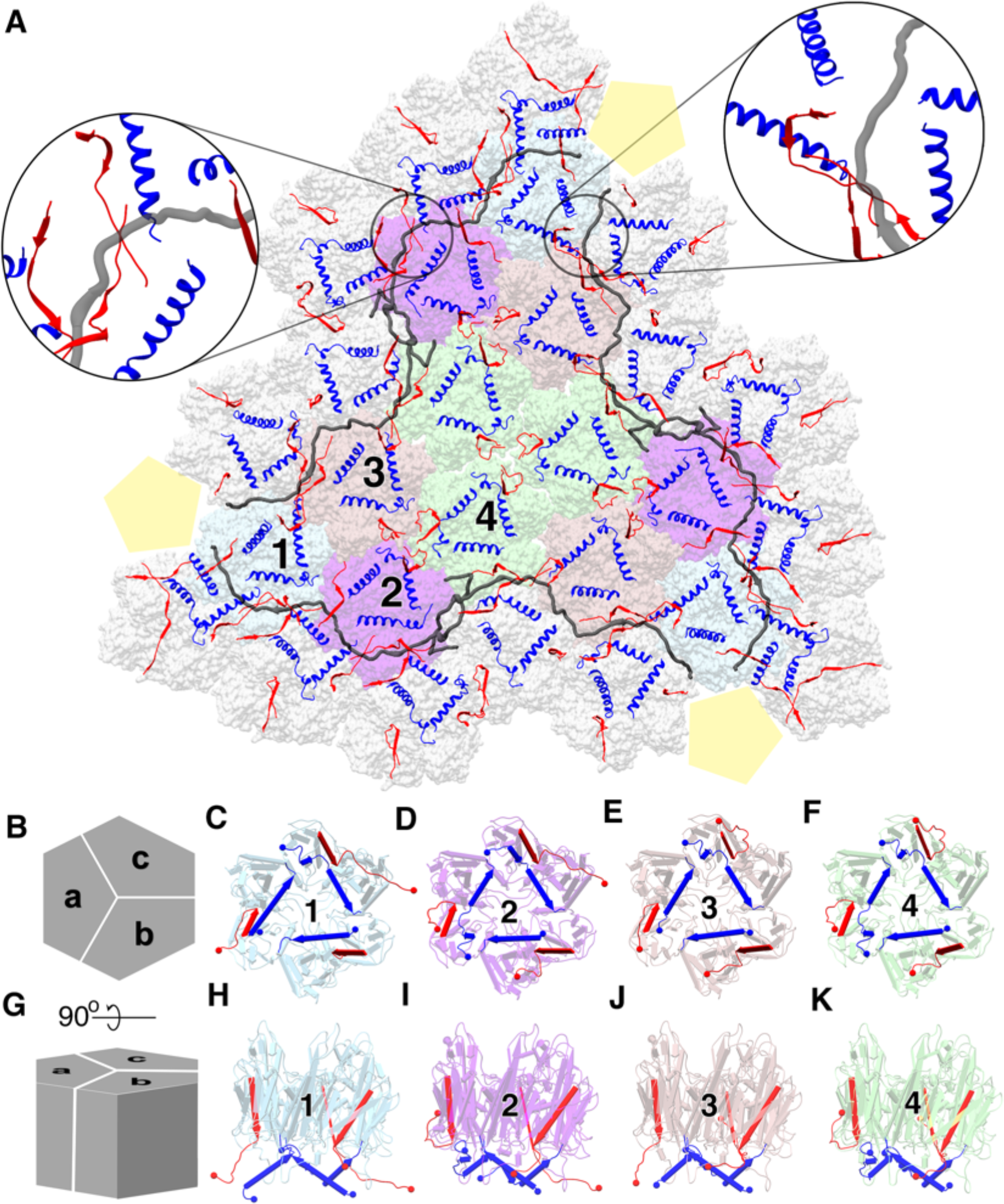
Major capsid protein P3 and its conformations. (**A**) (visualized from the outside of the viral particle) The 4 unique P3 trimers (represented in 4 different colors) and their arrangement forming the trisymmetron bound by the P30 dimers (in gray). The C-terminal region and the N-terminal region of P3 subunits are colored in red and blue respectively. The highlighted regions show the locking of P30 (in black) by the C-terminal region of the neighbouring P3 subunits, leading to the formation of a hinge-like mechanism which is not seen in PRD1. (**B**) Schematic represetation of the P3 trimers and the subunit arrangement to form a hexagonal capsomer (as viewed from outside) and (**C**), (**D**), (**E,** (**F**) are aligned to this view. (**G**) It is the orthogonal view to the schematic (**B**) and (**H**), (**I**), (**J**), (**K**) are aligned to this view. Different views of trimer 1 (**C,H**), trimer 2 (**D,I**), trimer 3 (**E,J**) and trimer 4 (**F,K**) show the variation in the N-terminal (shown as blue cylinders with the arrow heads pointing towards the C-terminal) and C-terminal (shown as red planks with the arrow heads pointing towards the C-terminal) region of P3 subunits. N and C termini are shown as spheres with respective colors. They are colored to match (**A**). The yellow pentagons are a schematic representation of the penton. *Also refer* Figure 2–figure supplement 1 and 2

**Figure 3:**
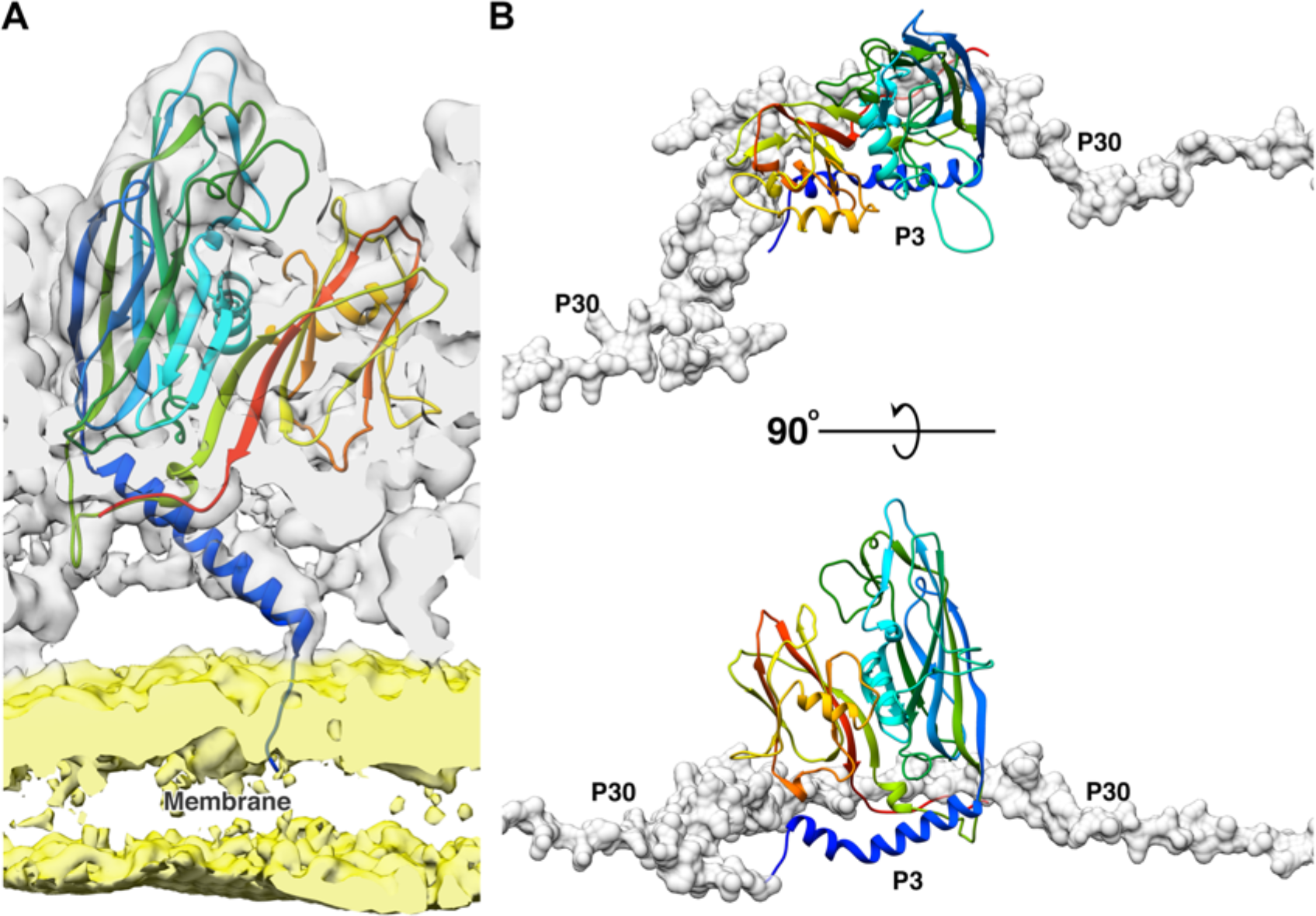
Function of 2 N-terminal helices of the P3 subunits. (**A**) Shows the model of the long N-terminal helix conformation (blue) of the P3 subunit anchoring to the membrane. The CryoEM map was low pass filtered to 5 Å to reduce noise in the membrane region due to map sharpening. In the CryoEM map, the capsid region is shown in grey and the membrane region is shown in yellow. (**B**) The newly discovered P3 N-terminal conformation, a long helix with a kink accomedates the P30 dimer (shown in gray, rendered as surface).

**Figure 4:**
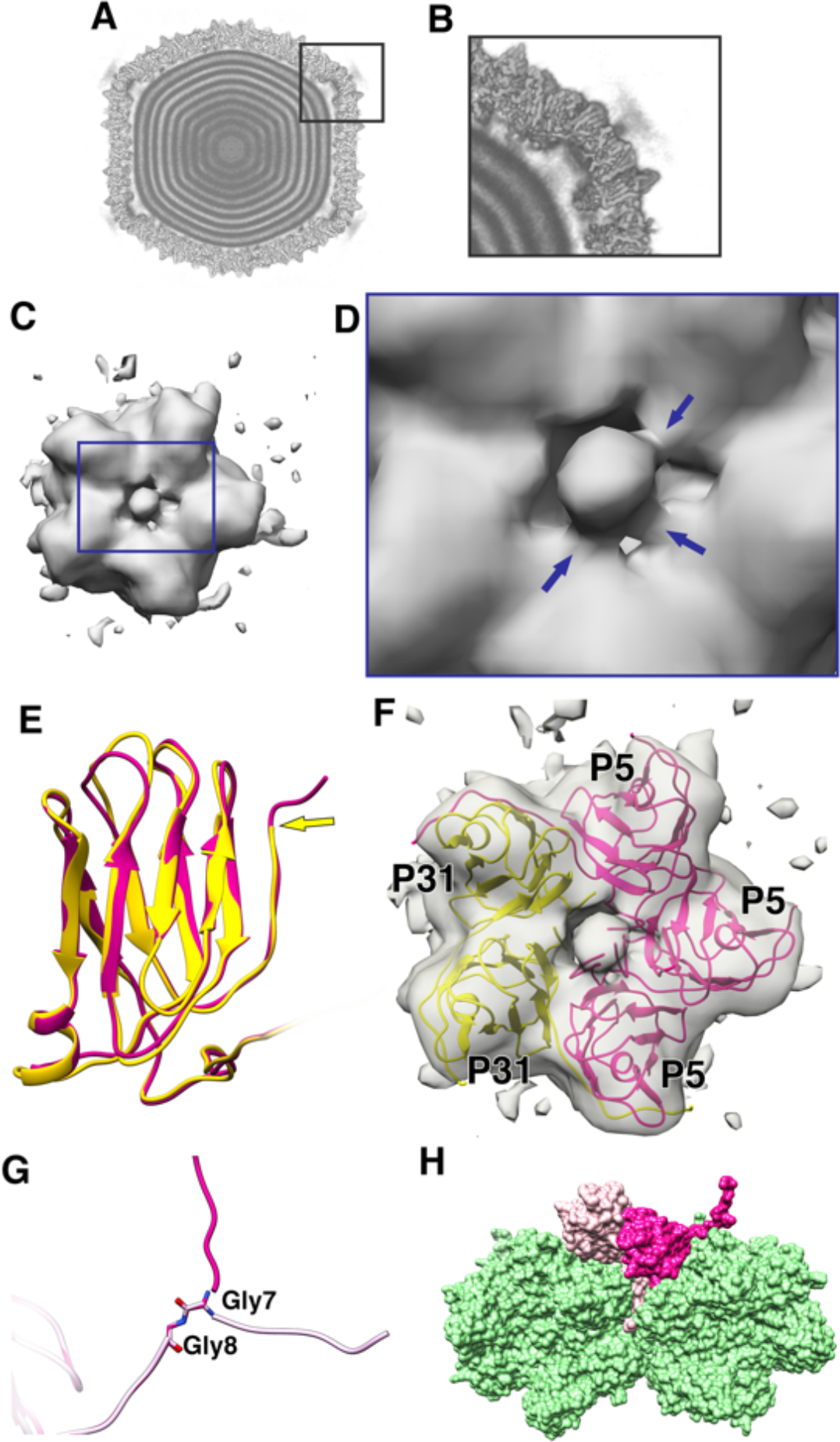
Penton as a heteropentamer of P5 and P31. **(A)** Icoshedrally averaged CryoEM map of PR772 showing the smeared densities at the five-fold vertices (Visualized using USCF Chimera, with volume viewer parameters: Style solid, step 1 and level 0.065, Plane 418, Axis Y, Depth 23). (**B**) Magnified image of a vertex showing the smearing of densities due to mismatch in symmetry. (**C**) Typical top view of the vertex map from the localized asymmetric reconstruction shown as a grey surface and the highlighted region is magnified in (**D**). In (**D**), the three stem-like protrusions (indicated by blue arrows) which interact with one another to form the stalk are shown. (**E**) The structure of P31 (yellow) and P5 (pink, residues 1-124) are superimposed and P31 terminates close to the five-fold (yellow arrow) and P5 (residues 121-124) continues upward. (**F**) The N-terminal domain of P5 and P31 are fitted into the localized reconstruction vertex map as rigid bodies and the P5 residues 121-124 fit into the stem-like density shown in (**D**). (**G**) Shows Gly7 and Gly8 residues at the N-terminal end of the P5 subunit and the two conformations (regular conformation as a bright pink structure and the special conformation as a pale pink structure). (**H**) Shows how the two conformations of the N-terminal region of P5 interact with the neighbouring protein. The regular confirmation of the N-terminal end of P5 (bright pink) hugs the neighbouring penton subunit whereas the special conformation (pale pink) is wedged in-between the P3 trimers (green). *Also refer* Figure 4–figure supplement 1-5

**Table 1.**
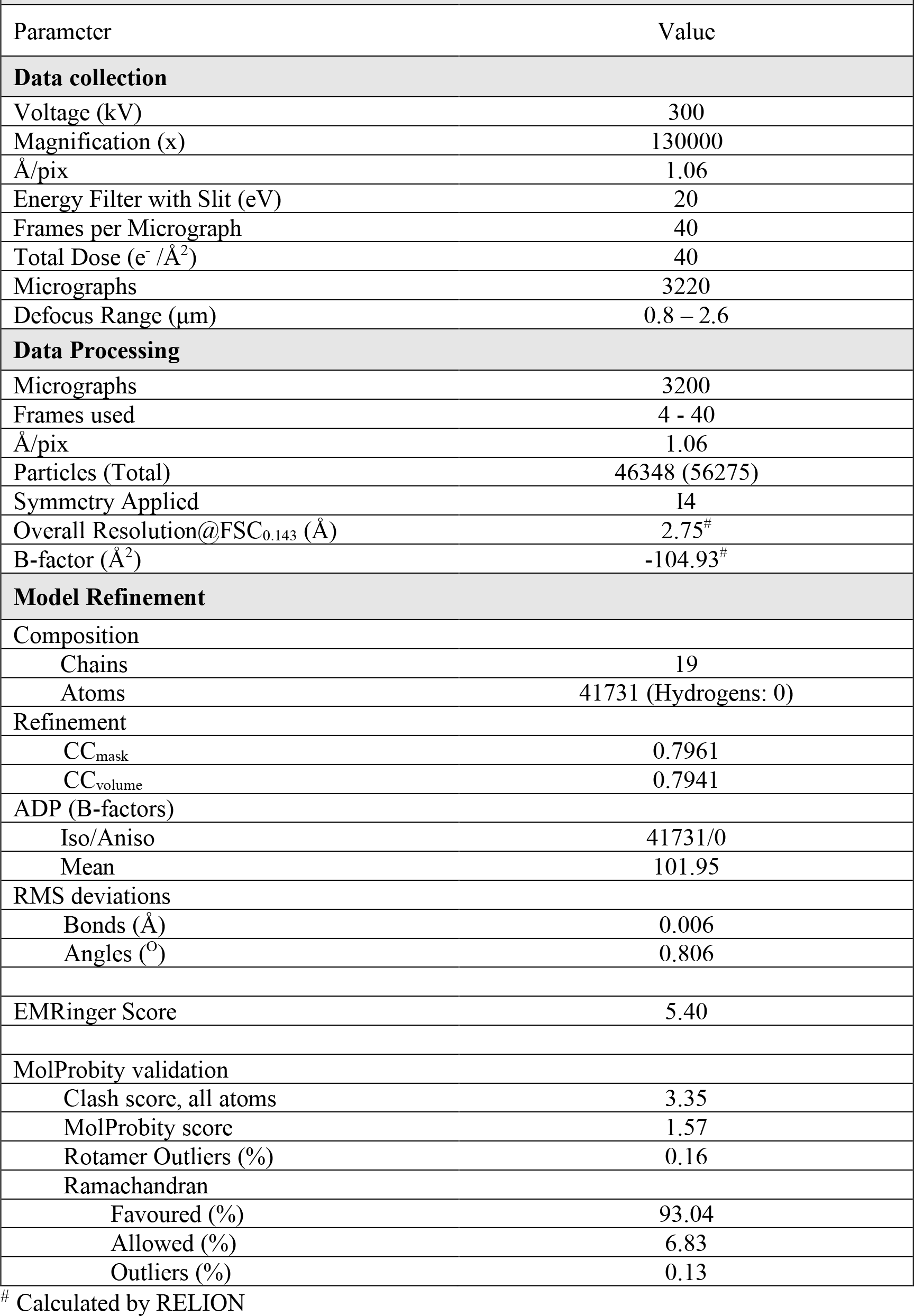
Data collection, Processing and Model refinement parameters.

The CryoEM 3D reconstruction of PR772 shows that the viral particle follows a pseudo T=25 lattice architecture with a (h,k) of (0,5) resulting in an icosahedral structure with 20 large trisymmetrons and 12 penta-symmetrons. However, the analysis revealed that the penta-symmetrons were hetero-pentamers (Caspar & Klug, 1962; Sinkovits & Baker, 2010). Each of these trisymmetrons have 36 copies of P3, the major capsid protein (MCP), arranged as 12 trimers, each of which structurally appears to be hexagonal in shape. At the fivefold vertices, the penta-symmetrons are replaced by a vertex complex to complete the icosahedral shell. With a pseudo T=25 architecture, PR772 is one of the larger wild type viruses resolved to a resolution below 3 Å.

### Major Capsid Protein and its conformations

P3 is the major capsid protein, which builds up the trisymmetrons of the capsid, and it is the most abundant protein found in bacteriophage PR772 (Figure 2A). Functionally, the P3 monomers (subunits a, b and c) are interlocked to form a trimer (Figure 2B-K). They exhibit the double-barrel trimer arrangement, as previously seen in viruses of the adenovirus linage (Benson, Bamford, Bamford, & Burnett, 1999; V. S. Reddy, Natchiar, Stewart, & Nemerow, 2010). The Leu130 - Ala150 loops from subunits a, b and c of P3 interact with each other in a cyclic fashion at the centre of the trimer complex to stabilise it. The monomers have a similar structure in bulk, with minor differences to accommodate the more significant variations in the C and N-terminal conformations. The asymmetric unit has 4 such unique trimers along with P30, P16 and a penton protein. Figures 2B-K show the four unique trimers and their subunit arrangement.

The N-terminal of the P3 monomers have 3 different conformations; a helix turn helix, a long helix and a long helix with a kink (Figure 2–figure supplement 1). Figure 2D is a good schematic to visualise all N-terminal conformations in a single trimer (subunit a shows helix-turn helix, subunit b shows long helix and subunit c shows helix with a kink). When the N-terminal adopts a helix turn helix, the shorter helix close to the N-terminal is bent away from the lipid membrane and interacts with the adjacent subunit of the trimer. The long helix with a kink behaves similar to a helix turn helix where the kink twists the helix away from the membrane but it is not embedded deeply into the adjacent subunit of the trimer. To accommodate the helix turn helix or the helix with a kink in the P3 subunits, the loop formed by residue Tyr351 – Val358 in these subunits is flipped. In case of the long helical conformation, the N-terminal residues Met1 – Gln6 anchor the P3 subunit to the lipid membrane (Figure 3A).

The N-terminal conformation of subunit a is more variable as compared to the conformation of subunits b or c in the 4 unique trimers. The N-terminal of subunit a can form either a helix turn helix bent away from the membrane or a long helix that anchors the subunit to the membrane. The N-terminal region of subunit c either forms a helix turn helix (Figure 2C, E, F, H, J, K) or a long helix with a kink (Figure 2D, I). The N-terminal of the b subunits always forms a long helix that anchors the subunit to the membrane (Figure 3). Subunit b of trimer 1, which is present close to the penton has the Tyr351 – Val358 loop flipped even though there is no helix turn helix or a long helix with a kink in this region that has to be accommodated.

The C-termini are more variable as compared to the N-termini of the trimers. They adopt 4 different conformations; one conformation is a long strand extending towards the lipid membrane and found only in subunit b of all trimers (Figure 2 C-F, H-K). The second conformation extends away from the membrane into the peripheral space between the trimers and it is also the most common C-terminal conformation. Two of the four instances in subunit a (Figure 2 D, F, I, K) and two of the four instances in subunit c (Figure 2 E, F, J, K) adopt this conformation. The third conformation is seen in subunit a of trimer 1, where the C-terminal runs parallel to the long N-terminal helix of the same subunit and it is embedded into the adjacent trimer 1 of the neighbouring trisymmetron (Figure 2 C, H). The fourth type of C-terminal conformation is seen in 2 instances of subunit c when they reside close to P30 (Figure 2 C, D, H, I). Here, the C-terminal is elongated and runs towards the lipid membrane, grazing it, while the N-terminal helix turn helix moves away from the membrane.

### Penton base is a heteropentamer of P5 and P31

The penton region of the icosahedrally symmetrized CryoEM map of PR772 was sectioned from the whole viral map using UCSF Chimera (Goddard, Huang, & Ferrin, 2007). To generate an initial model for de-novo model building of the penton region, PHENIX (Adams et al., 2010): Find Helices and Strands was used with both RESOLVE and PULCHAR options enabled. Due to the high sequence similarity between P31 and N-terminal P5 domain we could not rule out that any of these two proteins or a mixture could potentially form the penton base. Therefore, two initial models were generated. One model used the P31 protein sequence and the other used the P5 protein sequence as part of the input. In the initial observation of all the predicted segments of the two models, the P31 protein segment with residues 112-126 and the P5 protein segment with residues 107-121 occupied the same region of the sectioned map (Figure 1E, F). On closer inspection of the side chains from the two models and their fit into the CryoEM map densities in this region, the P5 protein side chains fit clearly at 3.24 RMS and the P31 side chains could only be fitted at RMS values lower than 2.64. As the CryoEM map was generated by icosahedral symmetrized reconstruction, we assumed that the densities of P5 and P31 were averaged. This hinted that the penton base could be a heteropentamer formed by both P5 and P31.

With an assumption that the penton base could be a heteropentamer, the P5 N-terminal domain (residues 1-124) and P31 (residues 1-126) were modelled using the icosahedrally averaged map at ∼3.04 and ∼2.2 RMS respectively in *Coot* (Emsley, Lohkamp, Scott, & Cowtan, 2010) (Figure 4E and Figure 4–figure supplement 3). The prominence of the amino acid side chain densities varied based on the protein sequence conservation between P5 and P31 (Figure 4–figure supplement 1). All identities and most of the conserved substitutions in the sequence alignment resulted in clearer well-defined amino acid side chain densities and the non-conserved substitutions resulted in amino acid side chain densities that were averaged between the respective side chain densities (Figure 1D-F and Figure 4–figure supplement 2).

Another unexpected feature that was observed at a lower RMS of about 1.04, a branching of the density close to the N-terminal region of either P31 or P5. One prominent branch, which was earlier used for modelling and another branch that is only visible at lower RMS. This led us to believe that the N-terminal of either the P31 or P5 could have an alternate conformation. On inspecting the residues from both the P31 model and the N-terminal domain model of P5, that were close to the branch region, P31 was ruled out as a potential candidate. P31 has Val11-Thr10-Met9 residues, which could not be fitted into the branched density without severely distorting the Cα backbone. P5 in the same region has Ser9-Gly8-Gly7. The 2 Glycine residues provided the needed backbone (Cα chain) flexibility that could facilitate this “special case” of N-terminal conformation (Figure 4G and Figure 4–figure supplement 5). In a typical arrangement, the N-terminal ends of both P5 and P31 would hug its neighbouring subunit counter-clockwise and stabilize the penton complex (Supplementary Figure 1). In the special case as described above, the N-terminal end of the P5 protein wedges itself between two adjacent P3 subunits (Figure 4H).

**Figure 5:**
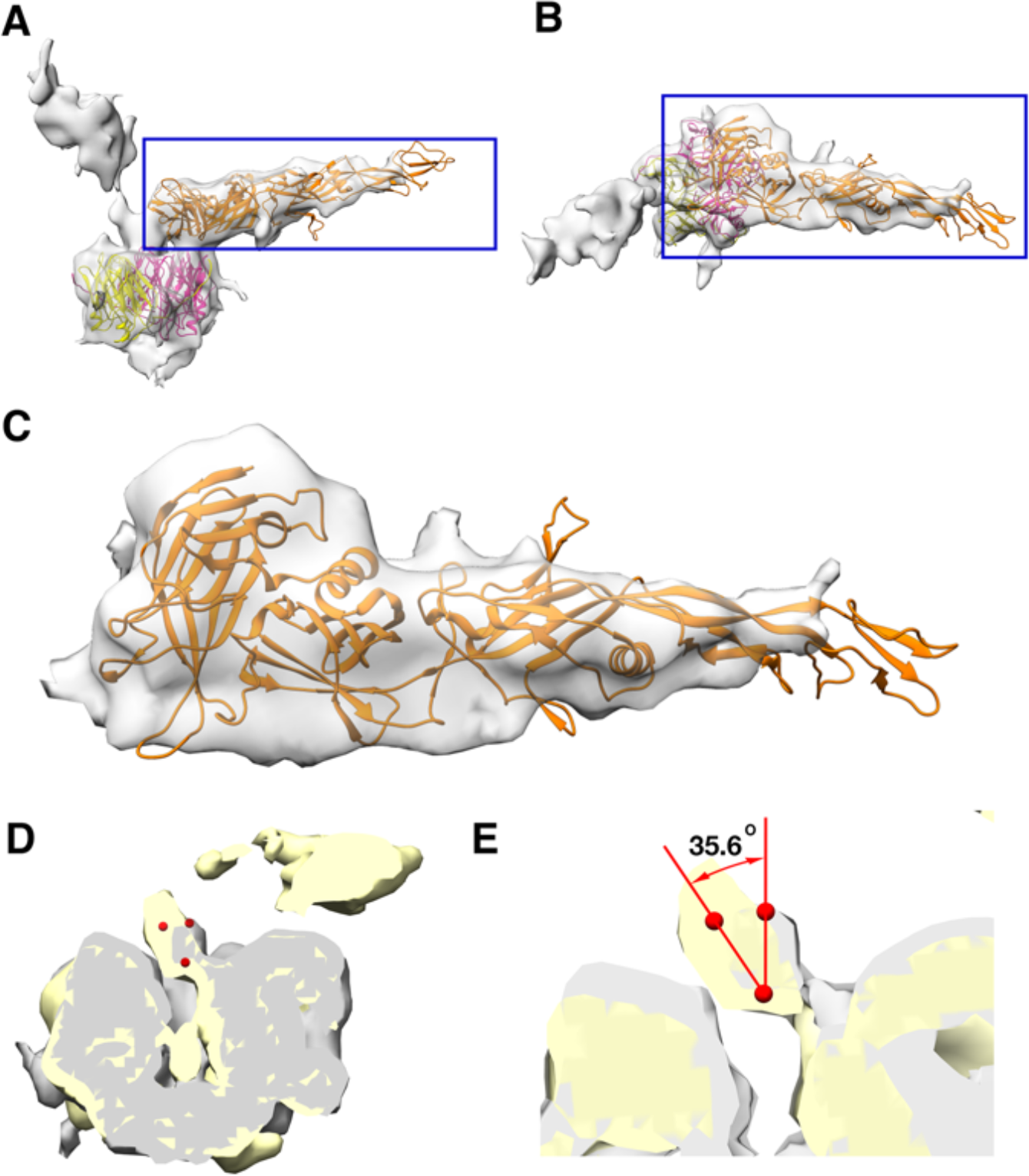
Monomeric P2 bound to P5. Localized asymmetric reconstruction of the vertex complex showing the two protruding densities as gray surfaces where (**A**) is the side view and (**B**) is the top view. P2 (orange), P5 N-terminal base (bright pink) and P31 (yellow) structures fitted into these map densities. (**C**) Isolated CryoEM density that represents the P2 subunit. It represents the region highlighted by the blue box in (**A**) and (**B**). (**D** and **E**) Superimposed vertex maps with P2 bound (yellow) and without P2 bound (gray). In classes where P2 is bound to P5, the stalk region is nudged by ≈35.6^0^ when compared to the classes where P2 is not bound to P5. Chimera: Volume Trace tool was used to place the red spheres in the density and Chimera: Measure Angles tool to determine the angles. *Also refer* Figure 5–figure supplement 1 and 2, Figure 5–movie supplement 1

**Figure 6:**
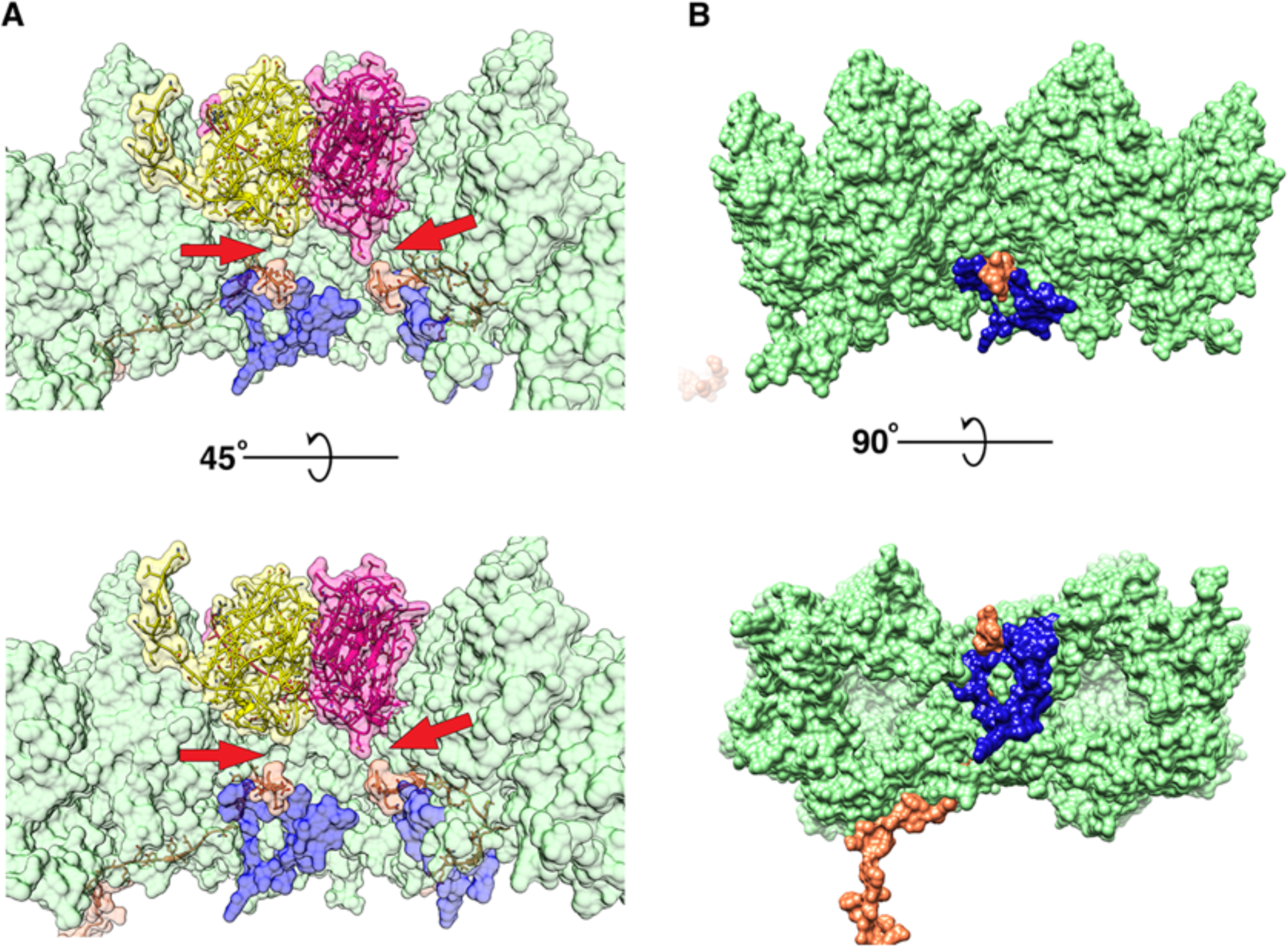
P16 and P30 intractions close to the vertex. **(A)** Shows the difference in interaction (pointed by red arrow) of the C-terminal Gly84 of P30 (orange) with P5 (bright pink), P31 (yellow) and P3 (green). The hydrophobic intraction of P30 with P5 is more obvious. (**B**) Shows the P3-P30-P16 complex (the view is similar to (**A**) but one copy of P16 and P30 are shown and the penton proteins are hidden), P16 (blue) lock the two adjacent P3 trimers (trimer 1)(green) and the P30 protein(orange). *Also refer* Figure 6–figure supplement 1, Figure 6–movie supplement 1

At 2.01 RMS, we noticed that the density for the C-terminal Gly84 of P30 was pointing towards the penton base and away from the inner lipid membrane. This is different from what is seen in bacteriophage PRD1 (Abrescia et al., 2004). In case of P5, the C-terminal Gly84 of P30 seems to interact with the Met19 residue of a P5 subunit by a hydrophobic interaction (Figure 6A). With P31, the hydrophobic interaction is not evident.

From the icosahedrally symmetrised reconstruction, at lower contour levels (0.065 in chimera) the map showed smeared densities above the 5-fold vertex of the viral particle (Figure 4A-B). This could be due to a symmetry mismatch of the proteins present in the region. Accordingly, the smeared region over the five-fold vertex was isolated and resolved by capsid signal subtraction followed by localized asymmetric (C1) reconstruction (see Methods). All the classes generated by 3D classification showed a single protruding density except one of the classes, which revealed two significant densities; one poorly resolved knob-like density and another more well resolved density closely interacting with one of the monomers of the penton (Figure 5–figure supplement 1). On closer inspection of every 3D class generated during the process of localized asymmetric reconstruction, we noticed that three of the subunits of the penton base had a stem-like protrusion close to the 5-fold axis, extending outwards and interacting with each other forming a thick stalk (Figure 4C-D). The other two subunits lacked the stem-like protrusion. The class that revealed the two significant densities also showed that one of the densities interacted with the thick stalk (Figure 5 A-B).

The number and arrangement of P5 and P31 forming the penton were confirmed by the localized asymmetric reconstruction. All the classes from the 3D classification showed that only three subunits formed the stem-like protrusion that interacted with one another to form a thicker stalk. P31 terminates close to the fivefold and thus cannot form the stem like protrusion whereas P5 residues (121-124) continue up and outward and these residues have the potential to form the stem like protrusion (Figure 4E and Figure 4–figure supplement 4). Three copies of the model of the P5 N-terminal base were fitted into the penton density from the localized asymmetric reconstruction. These models did indeed fit the density (Figure 4F). The overall map:model correlation reduced when the models of P31 and P5 N-terminal domain were swapped with each other in the localized reconstruction density map. The orientation and alignment of these residues also confirm previous predictions, which showed the formation of a triple helix with a collagen-like motif (residues 124 - 140) (Huiskonen et al., 2007; Javier Caldentey et al., 2000). The poorly resolved knob-like density represents the trimerized C-terminal domain of P5. By this it can be concluded that 3 copies of P5 and 2 copies of P31 form the penton base in PR772 (Figure 4F).

### P2 monomer is bound to P5 and stabilized by the P5 stalk

With the localized asymmetric reconstruction, the structure and composition of the penton base was revealed. It is now known that the stalk-like density observed, emanating from the penton base, is built with three copies of the P5 protein, forming a collagen-like motif. As described previously, one of the 3D classes, with ∼41,000 sub-particles from a total of 276,000 sub-particles, also revealed an extra rigid density closely interacting with the penton base and the protruding stalk (Figure 5 A-C and Figure 5–figure supplement 1).

In previous studies, it was shown that P2 and P5 could be interacting at the vertex (Bamford & Bamford, 2000). The crystal structure of P2 (PDB:1N7U) from PRD1 (98% protein sequence identity to PR772) was fitted into the density. The fit was good and the density that was resolved by localized reconstruction was sufficient to fit the domains I and II, forming the head and the domain III, forming the tail of P2. The off-centred arrangement of the tail with respect to the head allowed us to specifically assign the map density to the respective regions of the P2 protein (Figure 5 A-C).

The interaction of P5 with P2 is similar to a “ball and socket joint”, where the N-terminal base of P5 is the ball with domain I and domain II of P2 acting as a socket (Figure 5–movie supplement 1). The estimation of the electrostatic surface potential of both P2 and P5 shows good charge complementarity in and around the regions of interaction. These regions also have similar hydrophobicity (Kyte-Doolittle scale) (Figure 5–figure supplement 2 and Figure 5–movie supplement 1). P2 also interacts with the collagen-like motif of the P5 stalk, thus introducing some rigidity to the C-terminal of the P5 trimer. The presence of P2 nudges the P5 stalk by ∼36° from its usual position (Figure 5 D-E). The P5 stalk acts as a linchpin that locks the P2 molecule and restricts it from swivelling around the 5-fold vertex, thus stabilizing the complex.

The occupancy of P2 on the 5-fold vertices appears to be significantly lower in bacteriophage PR772 than what was observed from other members of the *Tectiviridae* family. The 3D classification of the signal subtracted and isolated vertices revealed that only ∼41,000 sub-particles from a total of ∼256,000 extracted sub-particles, showed the presence of a density representing P2. This accounts for 16% of all the sub-particles used or about 2 vertices in an intact viral particle, assuming an equal distribution among the viral particles.

### Overall Architecture of PR772

P30 adopts an extended conformation in an intact viral particle and it is found wedged in between the trisymmetrons (Figure 2A, S16 Fig). Two copies of P30, interlocked at the N-terminal hook, span between the adjacent vertices. In an intact viral particle, P30 forms a cage-like structure, which stabilises the trisymmetron and also interacts with the neighbouring trisymmetrons (Figure 2A and Supplementary Figure 2). The residues Tyr62 – Ile64 and Val32 – Arg35 of P30 form beta sheets with residues Thr384 – Leu386 from subunit c of P3 trimer 1 and residues Thr384 – Asn388 from subunit c of P3 trimer 2 respectively (Figure 2A highlighted regions). These C-terminal regions of P3 are sandwiched between P30 and the lipid membrane (Figure 2A highlighted regions). P30 is also sandwiched between two adjacent P3 subunits of the trimer from a neighbouring trisymmetron (Figure 2A, 6B and Figure 6–movie supplement 1).

Close to the penton, P16 forms a clamp-like complex using the loop Asn44 -Val55 and helix Asn101 - Ala115, which latches onto the loop Val242 - Tyr 247 of one of the P3 subunits of trimer 1 and locks it with the adjacent P3 subunit of trimer 1 from the neighbouring trisymmetrons. This further locks the P3-P30-P3 sandwich and makes it rigid. To accommodate the P30 protein in the P3-P30-P16 complex (Figure 6B and Figure 6–movie supplement 1) the loop formed by residues Tyr351 – Val358 in the P3 subunit in trimer 1 is flipped, as was mentioned earlier.

Three copies of P5 and two copies of P31 are held together by their N-terminal residues. Met9 – Val14 of P31 and Ser9 – Tyr13 of P5 inter-digit with the neighbouring subunit of the pentamer as β-sheets. Residues Gln8 – Asn2 and Gly8 – Met1 of P31 and P5 respectively continue further and hug the neighbouring subunits (Supplementary Figure 1). As was mentioned earlier, residues Gly8 - Met1 of P5 can in special cases have an alternate conformation where they are wedged in between the adjacent P3 subunits (Figure 4H). There is no obvious direct interaction between the penton and the P3 subunits of the capsid except for the special cases mentioned above. The disorganised region of P16 seems to interact with the penton base and helps in binding the vertex complex to the trisymmetrons. (Supplementary Figure 3)

The long C-terminal trans-membrane helix of P16 (Leu7 - Ala28) was poorly resolved and barely visible at 0.6 RMS. The high-resolution structure of the disorganized region (Tyr58 - Ile96) of P16 also evades us. It was poorly resolved with no visible side chains and part of the density map representing the Cα backbone was missing even at very low RMS. Due to poor signal to noise ratios, these regions were not modelled, but the difference density map of the vertex region and the modelled penton showed that the C-alpha backbone continued into the empty pocket beneath the penton and also interacted with the penton close to the 5-fold axes (Supplementary Figure 3).

## Discussion

The trisymmetron of PR772 is composed of 12 trimers of the P3 protein that are in turn bound by P30, an overall arrangement similar to that of the close relative PRD1. With PR772, there is noticeable variations in the structure of protein subunits when compared to PRD1 (Supplementary Text 1). Unlike in PRD1, the high-resolution map of PR772 reveals that the N-terminal region of P3 has 3 conformations and one can easily differentiate between a long helix with a kink and a helix turn helix. The long helix with a kink is structurally critical to accommodate the interlocking region formed by the N-terminal hooks of P30 during the initiation of particle assembly (Abrescia et al., 2004; Butcher et al., 2012) (Figure 2A and 3B). The helix turn helix motif of P3 interacts with the adjacent subunit of the trimer and stabilises the complex (Figure 2). The long helix conformation of P3 helps in anchoring the trimer complex to the membrane (Figure 2 and 3A). In conclusion, the N-terminal region of P3 shows plasticity and play an important role in both stabilising the trimer and anchoring the trimer complex to the membrane. The C-terminal region of P3 also shows differences in its conformation as compared to PRD1. In trimer 1 (Figure 2A, C and H) subunit a and subunit c have elongated C-terminal regions that interact with the adjacent P3 subunits of the neighbouring trisymmetrons and in-turn locking on to P30 (Figure 2 right hand side insert; and Figure 6). A similar arrangement can be seen in the C-terminal region from subunit c of trimer 2 and subunit a of trimer 3 of the neighbouring trisymmetrons (Figure 2 left hand side insert). The presence of P16 close to the penton region makes the interaction of all the neighbouring trimer 1 more rigid (Figure 6) and also anchors the whole vertex complex to the membrane (Figure 7), emphasising the role of P30 and P16 in maintaining the size and structure of the icosahedral viral capsid.

**Figure 7:**
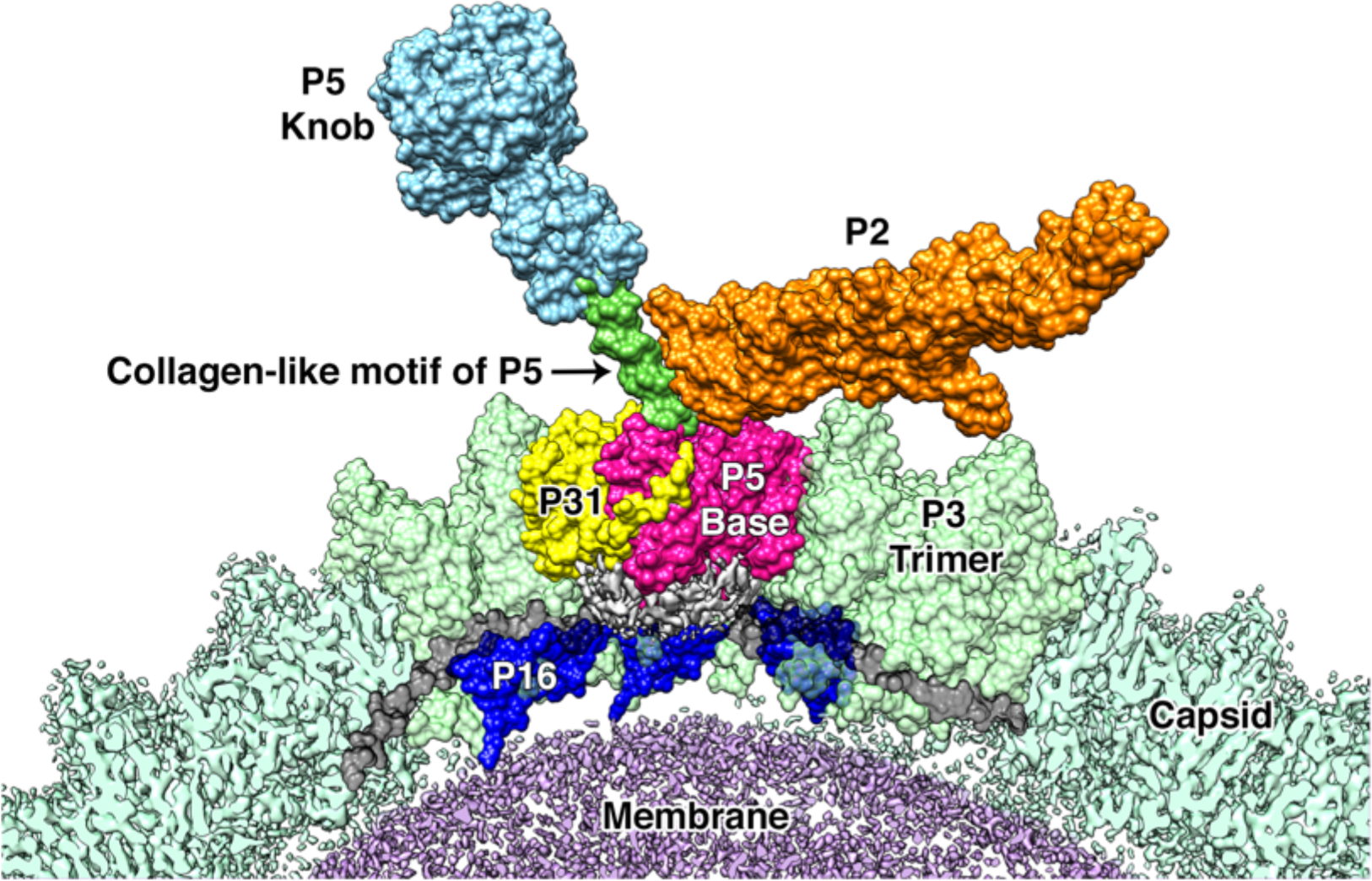
Proposed model for the vertex region. Shows the structural arrangement of P5 (P5 N-terminal base shown in bright pink, Collagen-like motif of P5 shown in bright green and P5 Knob is shown in pale blue), P31 (yellow) and P2 (orange) that form the vertex. The region below the vertex complex (shown as a grey density) is the unorganised/unmodeled region that links the surrounding P16 (blue) and the P30 (black). Capsid is shown in pale green (raw CryoEM map is labelled as Capsid and the modelled region is labelled as P3 Trimer). The lipid membrane is shown in purple.

Our studies show that unlike what was predicted for PRD1, the penton base of PR772 is an asymmetric heteropentamer consisting of three copies of P5 and two copies of P31 (Figure 4F). P31 has high sequence similarity with the N-terminal domain of P5. Previously, it was shown that P31 can replace P5 to form the penton and produce an intact viral particle (Bamford & Bamford, 2000). Even though the viral particles were intact, they were non-infectious due to the lack of the viral receptor-binding protein, P2, that is bound to the host recognition protein, P5 (Bamford & Bamford, 2000; A M Grahn, Caldentey, Bamford, & Bamford, 1999). In another study with the PRD1 sus525 mutant that lacks P31, it was also shown that these particles lacked the vertex complex (Rydman et al., 1999). With the current model, P5 will be unable to replace

P31 to form an intact viral particle due to obstructions that occur during the formation of the vertex complex. A penton with only P5 subunits will inhibit (i) the formation of a stable collagen-like motif (Figure 7) and (ii) the formation of the C-terminal host recognition domain of P5, as this structure only needs three copies of P5. A heteropentameric base of the vertex complex with three copies of P5 and two copies of P31 address both these issues.

The members of the *Tectiviridae* family have been shown to have structural similarities with adenoviruses despite infecting different hosts. In adenoviruses, the symmetry mismatch seen in the vertex region is solved by the tail region of the trimeric fiber interacting with three of the five grooves formed by the subunits of the penton base (Cao et al., 2012). Unlike adenoviruses, a similar problem in PR772 is solved by domain swapping. The trimeric P5 protein responsible for the formation of the spike, swaps the N-terminal base domain with the three copies of the P31 protein (Figure 4F).

The localized asymmetric reconstruction shows that P2 is bound to P5 (Figure 5). In the current CryoEM data, the orientation of P2 with respect to the penton base is reversed when compared to what was described for PRD1 (Huiskonen et al., 2007; Xu et al., 2003). Here, the beta-propeller motif of P2 with domain I and II, interacts with the N-terminal base of P5 and also the stalk. This was also speculated on in a previous study, where it was noted that members of the *Tectiviridae* family with large sequence variation in the beta-propeller region of P2 also showed variations in the P5 subunit as a compensatory effect (Saren et al., 2005). P2 interacts with both the N-terminal base and the stalk region of P5 but does not interact directly with any other structural protein in PR772. The interaction of P2 with the P5 stalk stabilises the C-terminal region of P5 (Figure 5). In the case of PRD1, P2 seems to occupy all vertices expect for the unique packaging vertex (Butcher et al., 2012; Javier Caldentey et al., 2000; Saren et al., 2005). In this scenario, the special conformation in which the N-terminal residues of P5 are wedged in between the neighbouring P3 subunits was found to be prominent (Abrescia et al., 2004). In the current model of PR772, P2 seems to occupy only ≈2 vertices of the icosahedral shell. This was also indicated in a previous study comparing the abundance of different viral proteins from various members of the *Tectiviridae* family using western blots (Saren et al., 2005).

In the icosahedrally averaged model of PR772, we see that the special conformation of P5 where the N-terminal residues are wedged between the P3 subunits of the adjacent trimer 1 is not prominent, but these P5 densities are visible at lower RMS values. The presence of P2 in the vertex complex may coincide with the N-terminal region of P5 adopting the special conformation. It was reported that, compared to the CryoEM map of a wild type PRD1 virion, the CryoEM map of *sus690*, a mutant PRD1 virion that lacks P2 and P5, showed absence of density in the region where we see the N-terminal of P5 wedged between two neighbouring P3 subunits. The lack of this density was attributed to the conformational changes in P31 due to the binding of P5 or the presence of P5 in between P31 and the P3 trimer (Huiskonen et al., 2007). In light of the current findings in PR772, P5 is known to be part of the heteropentameric base forming the penton and the absence of the above-mentioned density confirms that P5 adopts the special N-terminal conformation and not P31.

The penton has no obvious direct interaction with the neighbouring P3 subunits of the capsid, except in the rarely occurring special conformation of P5 where the N-terminus is wedged in between P3 subunits of trimer 1 as mentioned above. The difference density map of the vertex region and the modelled penton shows that the disordered region of P16 interacts with the penton. These interactions of P16 with P3, P5, P31 along with the membrane seem to play a central role in the formation of the vertex complex and anchoring it to the membrane.

From the localized asymmetric reconstruction of the vertex complex, one can notice that there is a variation in the map density below the penton. The map density below the P5 subunits is significantly different when compared to the same region below the P31 subunits. This suggests that, along with the interactions of P16 with P5 and P31, there could also be poorly resolved proteins like P11, P7, P14 or P18, etc; in the region (Luo, Butcher, & Bamford, 1993; Mattila, Oksanen, & Bamford, 2015; Rydman & Bamford, 2000). The variation in the map density in this region is also compounded by other interactions, like that of the C-terminal Gly84 of P30 with the Met19 of P5 subunits and lack thereof with respect to the P31 subunits.

Preliminary results for the whole particle asymmetric reconstruction using symmetry relaxation in EMAN2 of the wild type PR772 do not show the presence of a unique packaging portal in the dormant particle in contrast to PRD1. All the vertices of PR772 show the heteropentameric arrangement of the penton with 3 subunits showing the stalk and the other two without the stalk (Figure 4 C-D, Supplementary Figure 4).

Figure 7 shows our model of the vertex complex in PR772. Using this model, we propose a mechanism for the initiation of infection. In analogy to the studies on PRD1, It is known that P5 is needed for host recognition, but the binding of P5 to the host is transient (A M Grahn et al., 1999). The host binding is stabilised by the high-affinity interaction of P2 and its receptor, locking the viral particle to the host (A M Grahn et al., 1999). The relative changes between P5 and P2, triggers the disruption of the vertex complex by pulling the N-terminal region of the P5 trimers that are wedged between the P3 subunits. This will in turn disrupt the interaction between the P5 proteins and P30 as seen in Figure 6. This disruption cascades further and disturbs the interaction between P30 and P16, rendering the whole vertex complex unstable. P30 dimers could also transduce these effects to the neighbouring vertices. The disruption of the vertex complex exposes the viral membrane and membranous proteins like P18, P11/P7, P32 etc; to the host surface to facilitate the formation of the membranous tube for DNA transport. P16, that holds the vertex complex, along with other membranous proteins, could act as a protein tether that would help in moving the lipid membrane closer to the host (Supplementary Figure 5).

## Materials and Methods

### Preparation and purification of PR772

Bacteriophage PR772 (ATCC® BAA-769-B1) was propagated on *Escherichia coli* K12 J53-1. It was purified as previously described (H. K. N. Reddy et al., 2017) and further concentrated to facilitate testing various concentration of the viral sample during grid optimization for CryoEM. This method yielded about 2-4 mL of viral particles with a concentration of 1 mg/mL using 10-20 agar plates. The sample was further concentrated to 20 mg/mL by using an ultracentrifuge. The sample was added into an ultracentrifuge tube and then a solution of Caesium Chloride in buffer (HEPES 20 mM, NaCl 100 mM, MgSO_4_ 1 mM, EDTA 1 mM, pH 8.0) at a density of 1.34 g/mL (g/cm^-3^) was gently layered on top. The mixture was centrifuged at 100,000 ×g for 30 mins. The intact viral particles migrated to the top as a fine band and the broken particles along with any free DNA that was released from the broken particles stayed at the bottom of the tube. The top band with the intact viral particles was extracted using a needle and syringe. The concentrated sample was dialyzed over-night with the above-mentioned buffer to remove caesium chloride.

### CryoEM Grid Preparation and Data Collection

The condition for CryoEM grid preparation was optimised for collecting a large number of particle images. For vitrification of the viral sample by plunge freezing into liquid ethane, we used a Vitrobot Mark IV (ThermoFisher). The best grid condition with uniform sample distribution was obtained by applying 3 μL of 7 mg/mL concentrated viral sample solution on a glow-discharged C-Flat grid CF-2/2-2C under 100 % humidity at room temperature. The data were collected on a Titan KRIOS (ThermoFisher) equipped with a K2 Summit (Gatan) direct electron detector and a GIF Quantum LS (Gatan) energy filter. All the data were collected at a magnification of 130 k in EFTEM mode with a pixel size of 1.06 Å. The slit width of the energy filter was 20 eV. The dose rate was 4.4 e^-^ per Å^2^ per second with a total exposure of 9 seconds resulting in a total dose of ∼40 e^-^/Å^2^. The total dose was distributed over 40 frames in each movie. 3220 movies were collected.

### Image Processing

The movie frames were corrected for beam induced sample motion and aligned using MotionCor2 (Zheng et al., 2017). The first 3 frames of the movies were skipped and the rest were aligned. These aligned frames were averaged with and without dose weighting. The non-dose weighted image stacks were used to estimate defocus and correct CTF using CTFFIND4 (Rohou & Grigorieff, 2015). All estimated fits of defocus and CTF were visually inspected. All images with significant astigmatism or a prominent ring due to crystalline ice around 3-4 Å were discarded.

### Whole Particle Reconstruction

A total of ∼3200 images were used to auto pick 56275 particles using template matching in RELION (Scheres, 2012) (version 2.1 beta 1) (Kimanius, Forsberg, Scheres, & Lindahl, 2016). The 2D classes generated by 2D classification of 710 manually picked particles were used as templates for auto-picking. The auto-picked particles were binned 2× during the extraction (box size of 429×429 and 2.12 Å/pix). Extensive reference free 2D classification was performed to remove any particle images with ethane contaminants or broken/empty viral particles. The classes with good 2D averages were selected and the particles from these classes were extracted. This resulted in 51893 particles that were used for 3D classification. RELION: 3D initial model tool, which is based on stochastic gradient descent, was used to generate an ab-initio reference map for 3D classification.The 2× binned particle images were used to generate this low-resolution icosahedrally averaged map. This low-resolution map was used as a reference for 3D classification. The 3D classification was performed with icosahedral symmetry (I4) applied. The most dominant class, with 46348 particles, was selected and the particles were extracted for final refinement.

Icosahedral symmetry (I4) was also applied during the final refinement. The refinement with the 2× binned particles reached Nyquist sampling (∼4.24 Å here). The particles from the final iteration step of the refinement were re-extracted without binning and further refined. The reference map was also scaled to match the new box and pixel size (858×858 and 1.06 Å/pix respectively) using e2proc3d.py from EMAN2.1 package (Bell, Chen, Baldwin, & Ludtke, 2016) (Supplementary Figure 6).

After refinement, the maps were corrected for Ewald sphere effects using RELION 3.0 beta 2. These maps were post-processed. A soft binary mask was generated using the 15 Å low pass filtered map extended by 10 pixels and with a soft edge of 15 pixels. An initial binarization threshold of 0.001 was used to include all the map features (i.e, internal membrane, etc) in the mask. This was used as a solvent mask during post processing. The map was corrected for the detector’s Modulation Transfer Function (MTF) and sharpened with an inverse B-factor. Overall gold standard FSC@0.143 was estimated using two independently processed half maps. ResMap (Kucukelbir et al., 2014) was used to estimate the variation of resolution across the two unfiltered half maps.

### Localized asymmetric reconstruction of the 5 fold vertex

The 5-fold region of the viral particle was reconstructed with capsid signal subtraction followed by localized reconstruction(Ilca et al., 2015) with relaxed symmetry (icosahedral to C1). Particles from the final iteration of the 2x binned reconstruction were used. Signal subtraction was performed to remove the contribution of the viral capsid and the genome to the map density. About 276,000 sub-particles of 100 pixels box size were extracted (from one half map) and used for 3D classification without image alignment. No symmetry (i.e, C1) was applied during the classification. The initial model was produced by back projecting the extracted sub-particles using the previously calculated Euler angles from the icosahedral averaged 3D refinement. The initial model was low pass filtered to 45 Å and this map was used as a reference model during 3D classification. The resolution of the expectation step was limited to 10 Å and a spherical mask of diameter 200 Å was added using the mask diameter and flatten solvent options provided by RELION to reduce the effects of systematic noise due to signal subtraction. The classes generated were analysed and the particles from the class showing two protruding densities were selected and the 3D classification was now further refined with parameters similar to what was mentioned above but with the local image alignment enabled. The part of the map density that we were interested in, was clipped due to the spherical mask that was used, so the particles were re-extracted with a larger box size of 140 pixels. These newly extracted particles with a larger box were back projected using the Euler angles calculated during the previous 3D classification and the resolution was limited to 10 Å in the expectation step.

### Model Building and Refinement

Modelling the whole particle was difficult owing to the large size of the particle and the limited RAM available. The whole viral map was sectioned using the sub-region selection option in Chimera. The sectioned maps were optimised by local sharpening using PHENIX: Autosharpen (Terwilliger, Sobolev, Afonine, & Adams, 2018). These sectioned maps were used to generate a crude model using PHENIX: find helix and loops (Adams et al., 2010). The crude model was further used for de-novo modelling of the proteins in *Coot* (Emsley et al., 2010). These models were further refined using PHENIX: Real space refinement to improve the model. All the models were individually refined and put together to form the asymmetric model. The model of the asymmetric unit was further refined against a new map encasing the asymmetric unit using PHENIX-Real space refinement. The model of the asymmetric unit was validated with *MolProbity* (Chen et al., 2010), Mtriage (Afonine et al., 2018) (Supplementary Text 2) and EMRinger (Barad et al., 2015).

## Acknowledgments

We would like to thank Kazuyoshi Murata and Naoyuki Miyazaki from National Institute for Physiological Sciences, Japan; for providing us the sample cryoem data set to test the feasibility of the project. We would like to thank Sjors Scheres MRC Laboratory of Molecular Biology, UK and Björn Forsberg from Stockholm University, Sweden for their valuable feedback in handling large CryoEM maps in RELION.

## Data Availability

CryoEM Density maps and atomic models that support the findings of this study have been deposited in the Electron Microscopy Database and the Protein Databank with the accession codes EMD-4461 (Whole particle reconstruction), EMD-4462 (Vertex Complex) and PDB ID 6Q5U (Atomic model of the asymmetric unit).

## Figure Supplements

**Figure 1–figure supplement 1.**
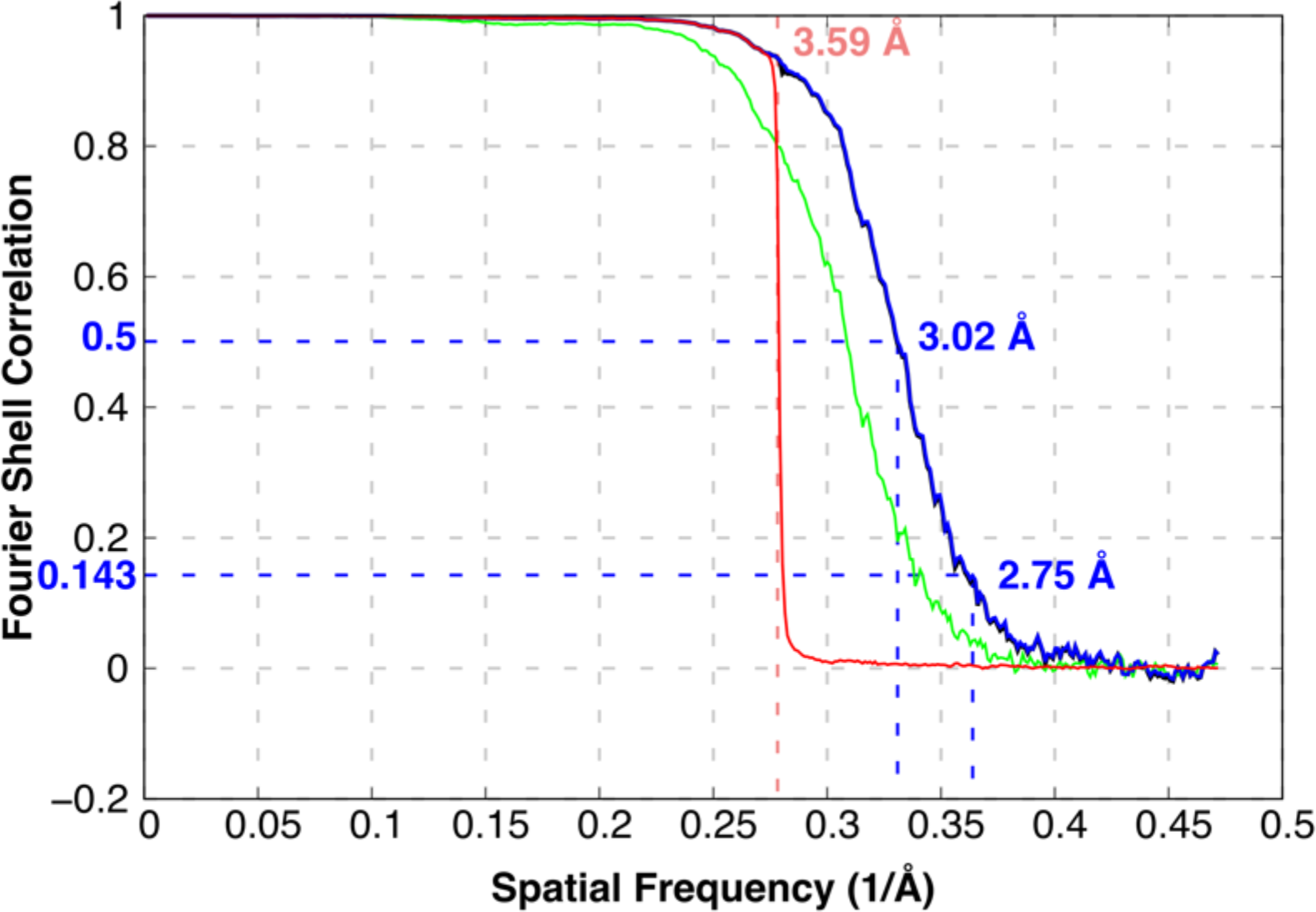
FSC curves of the overall resolution estimate of the 3D reconstruction (Green: unmasked map, Blue: masked map, Black: FSC corrected, Red: phase randomized map). FSC@0.143 is 2.75 Å and the FSC@0.5 is 3.02 Å. The phase randomization to test model bias was done at 3.59 Å. *Also refer **Source Data File 1***

**Figure 2–figure supplement 1.**
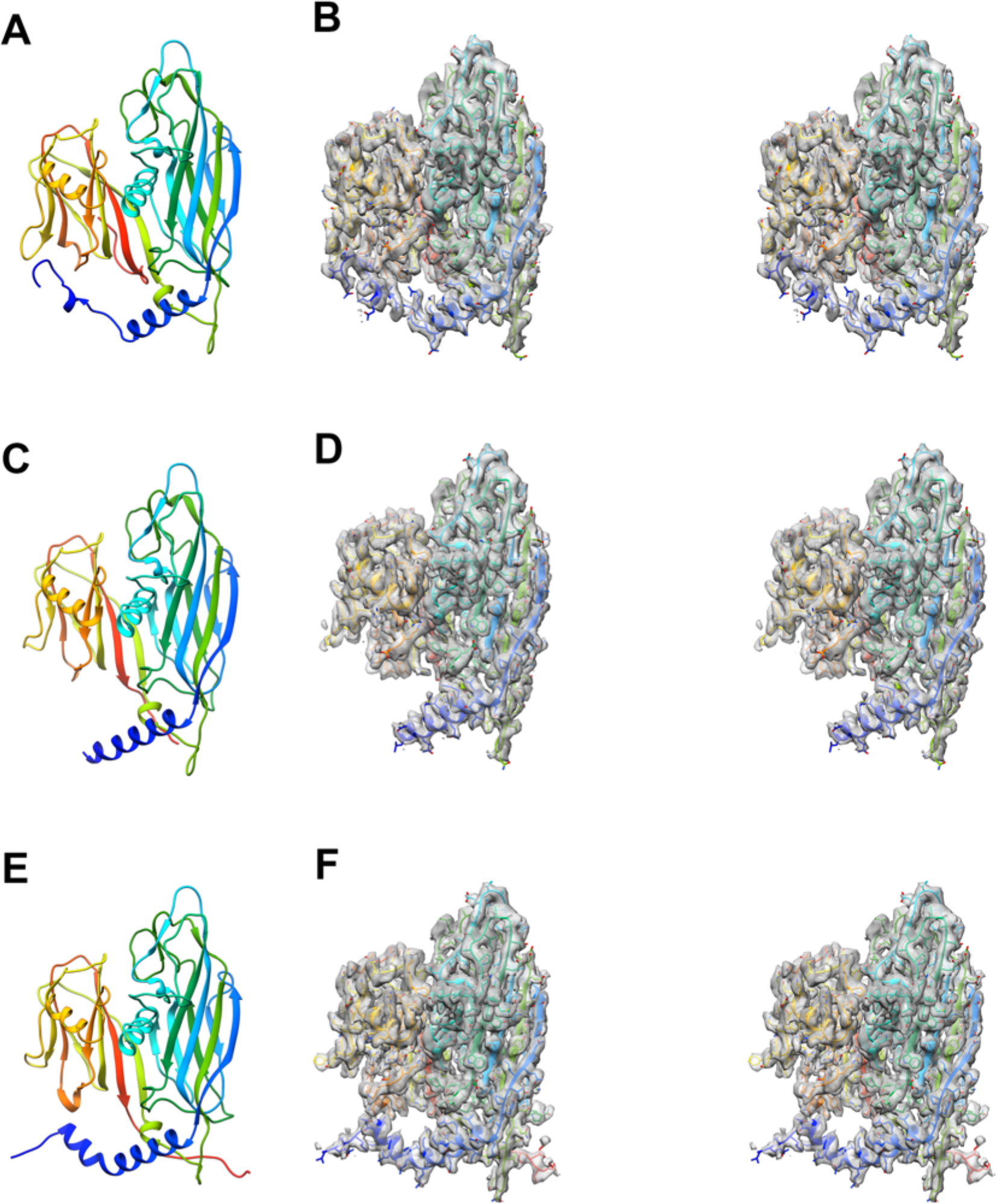
Shows the different N-terminal conformations of the P3 monomer. **A, C** and **E** represents the model of subunits a, b and c respectively of the trimer labelled 2 in Figure 2. **B, D** and **F** represents the stereo images of the model of subunit a, b and c fit into their respective CryoEM maps. The N-terminal conformations of P3 in **A, C** and **E** represent the helix turn helix, the long helix and the long helix with a kink respectively.

**Figure 2–figure supplement 2.**
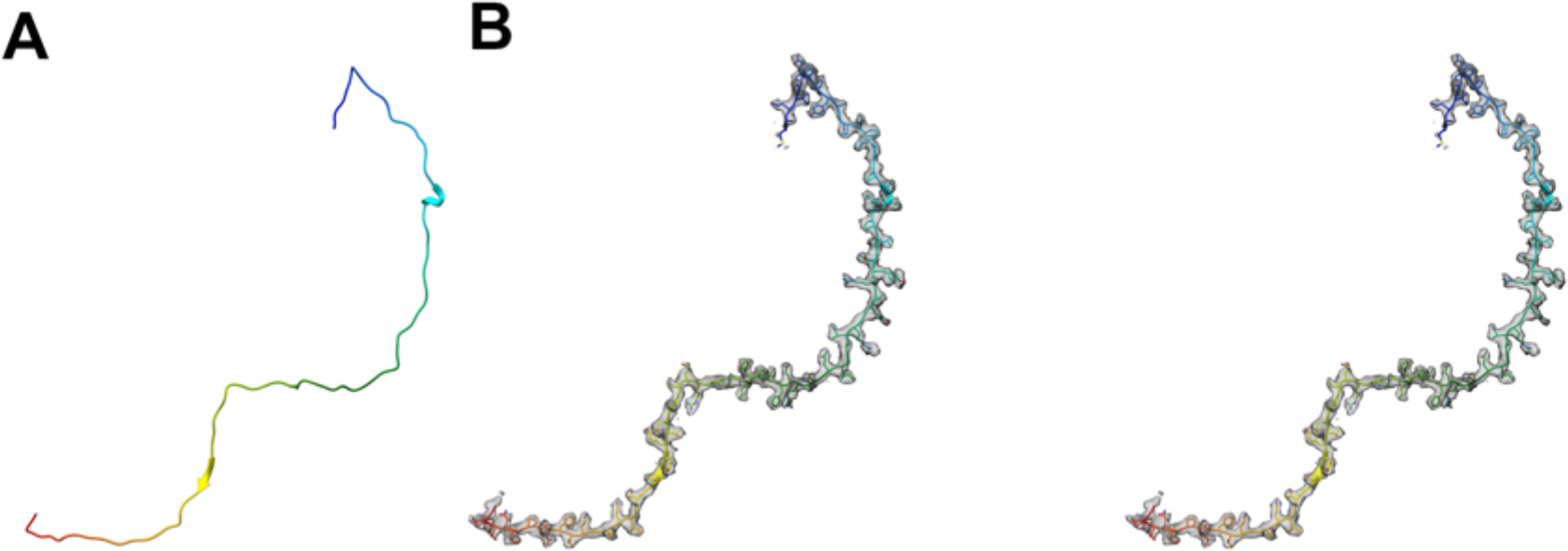
(**A**) The model of the tape-like protein, P30. (**B**) The stereo images of the model fitted into the CryoEM map.

**Figure 4–figure supplement 1.**
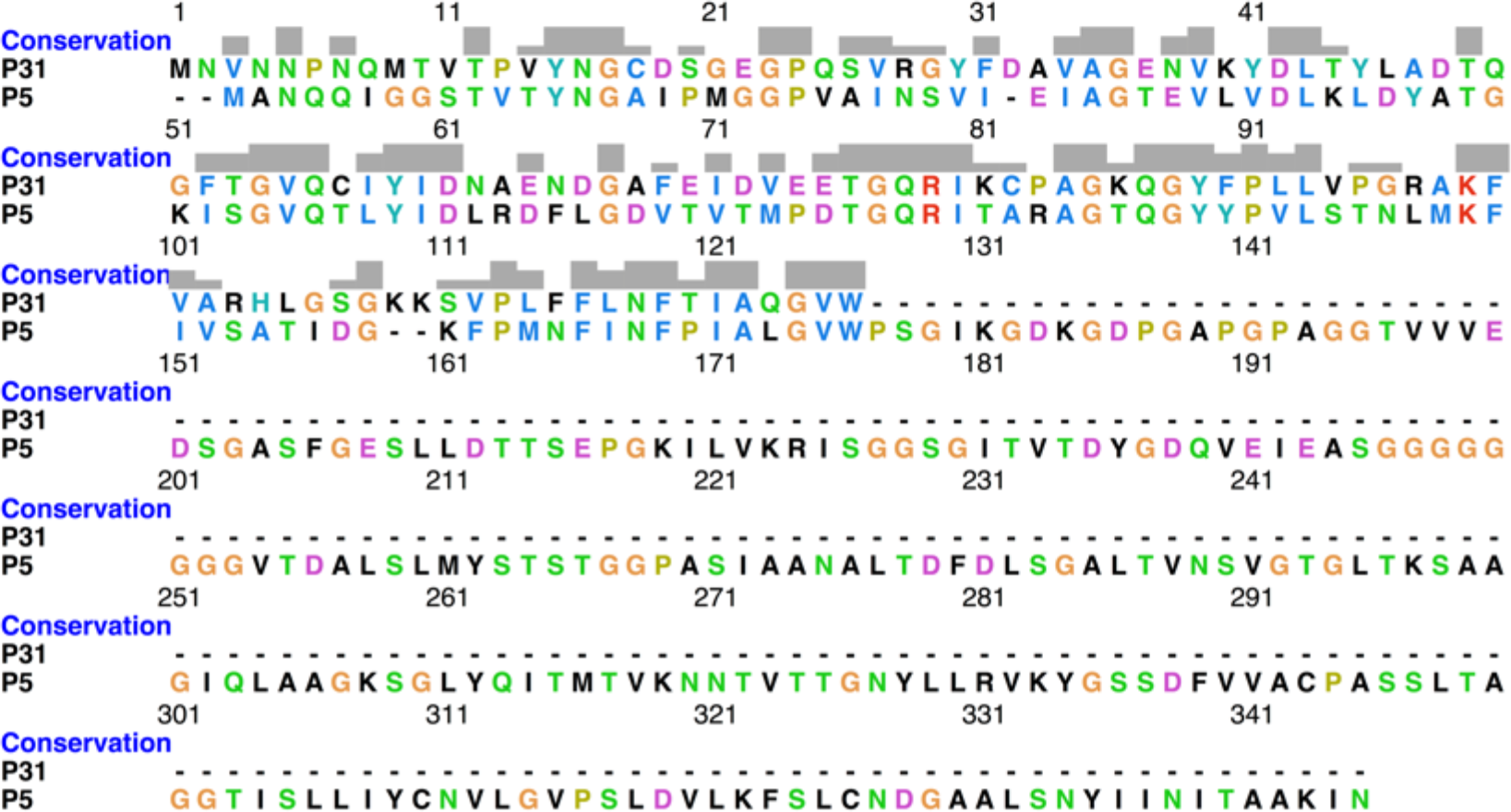
Shows the protein sequence conservation between P31 and P5. The protein sequence of N-terminal domain of P5 and P31 show 38.3 % identity and 71.3% similarity.

**Figure 4–figure supplement 2.**
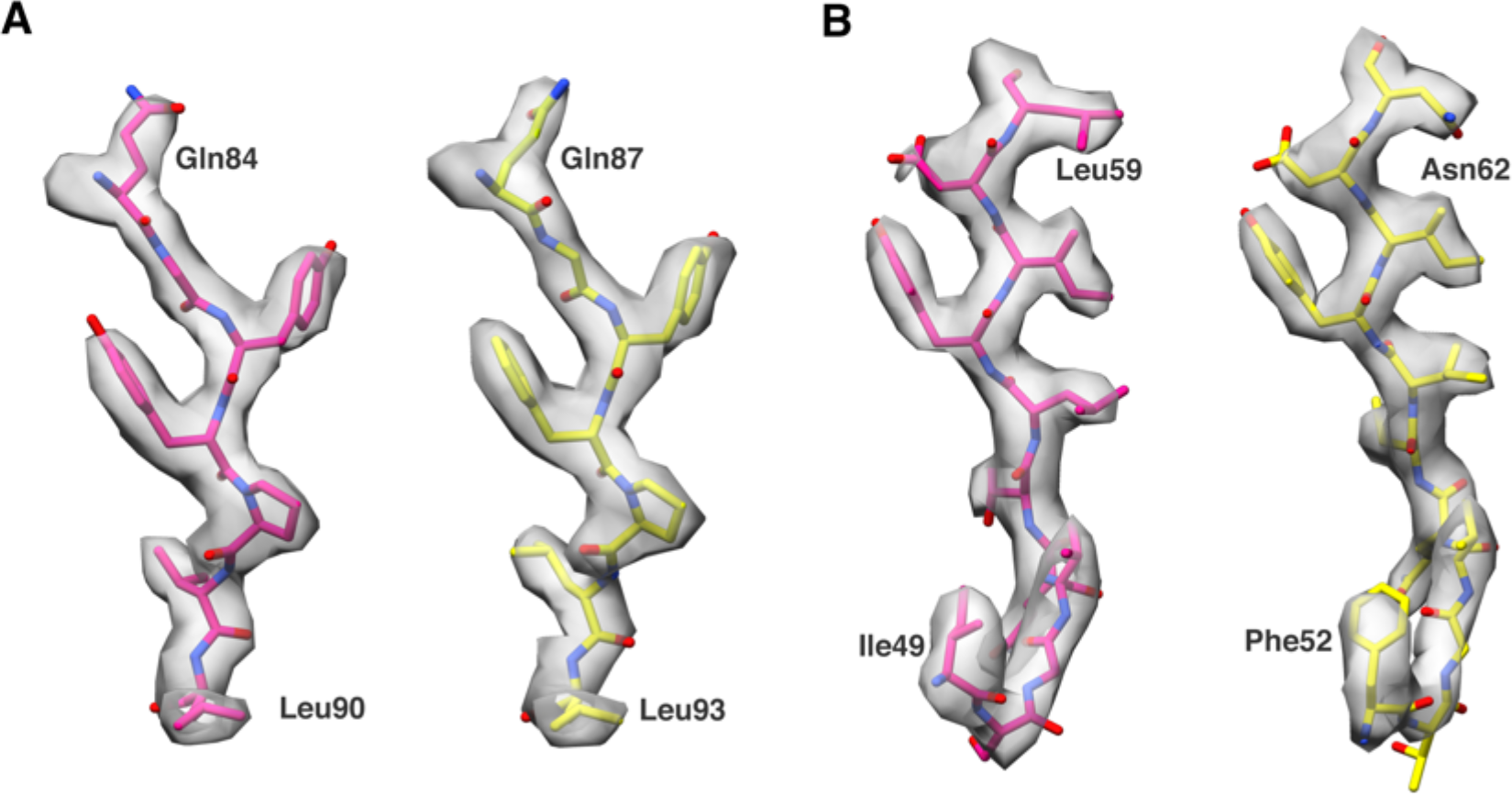
Comparison of model fitted into map density in few of the conserved regions between P31 and P5. P5 and P31 models are shown in pink and yellow respectively.

**Figure 4–figure supplement 3.**
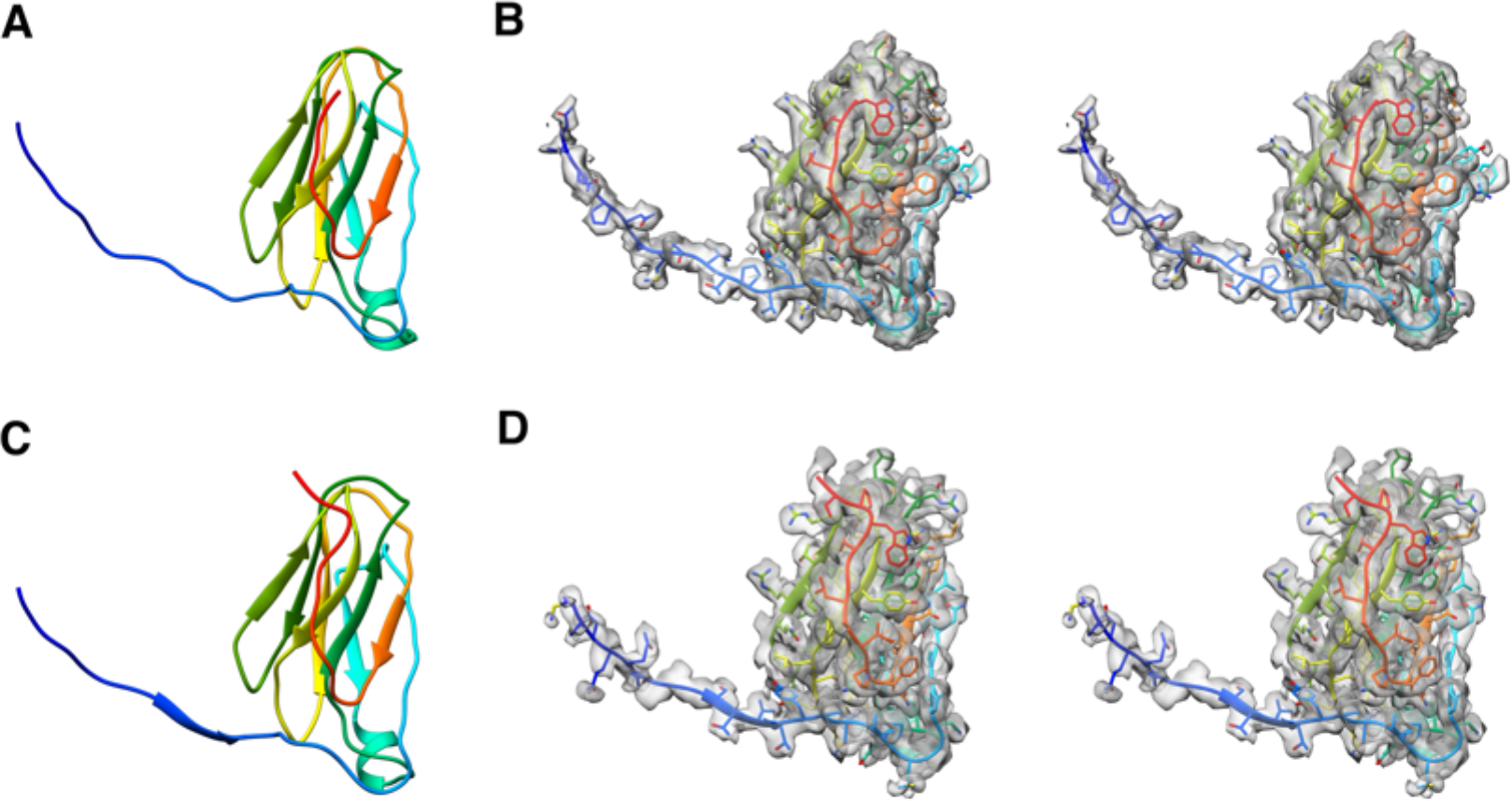
(**A)** and **(C**) are the models of P31 and the N-terminal domain of P5 respectively. (**B**) and (**D**) are the stereo images of P31 and the N-terminal domain of P5 fitted into the CryoEM map respectively.

**Figure 4–figure supplement 4.**
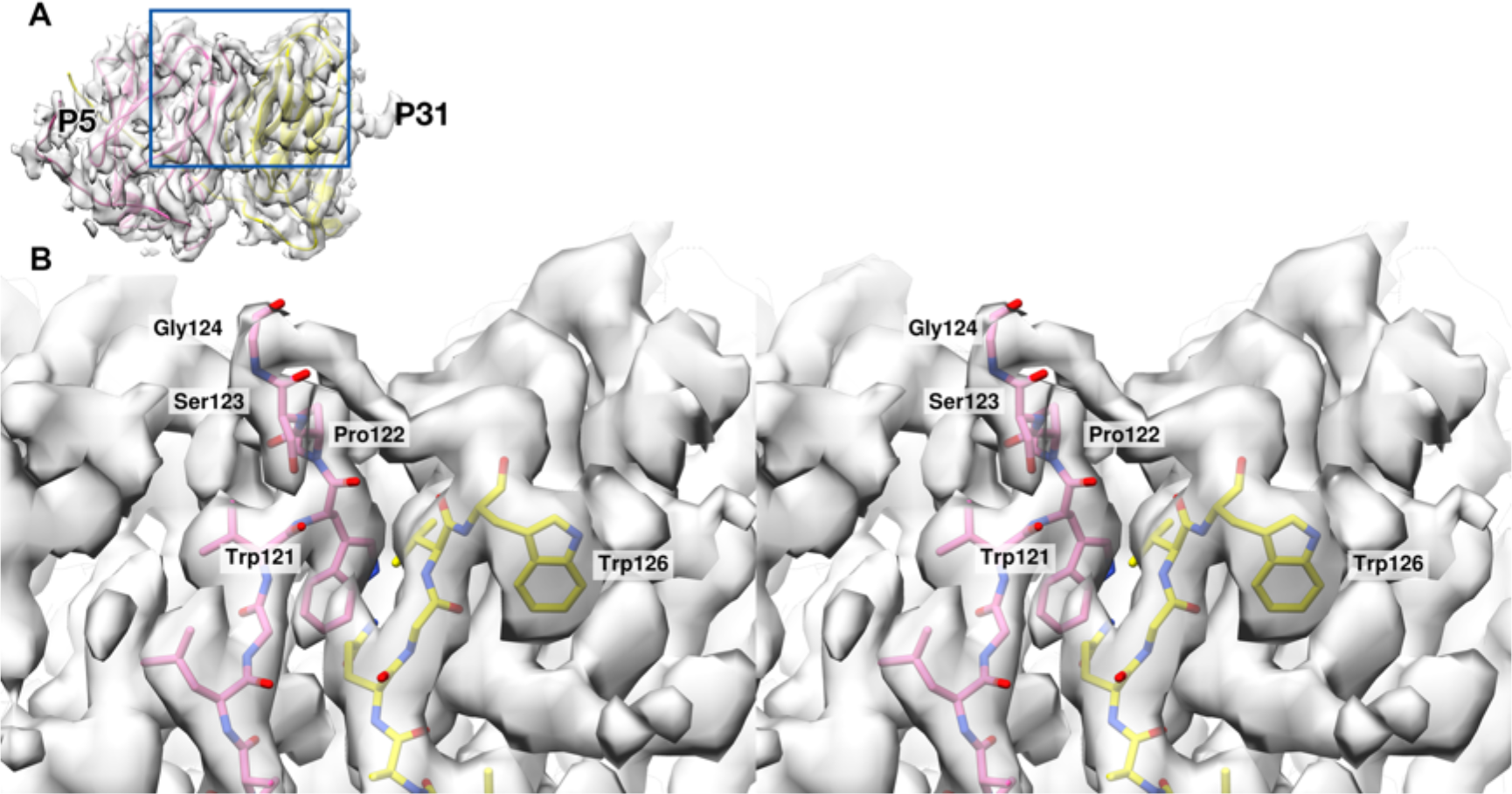
Comparison of residues from the N-terminal domain of P5 that contribute to the formation of the stalk and the lack of residues in P31 that contribute to the density. (**A**) shows the overview of the map:model for the adjacent subunits of P31(yellow) and N-terminal domain of P5 (pink). Blue box highlights the region shown in (**B**). (**B**) shows the stereo view of the region highlighted in (**A**). The P5 residues Trp121 - Gly124 (shown as pink sticks) extend upwards filling the map density, but there is no contribution from P31 (shown as yellow sticks). Trp126 of P31 is the terminal residue.

**Figure 4–figure supplement 5.**
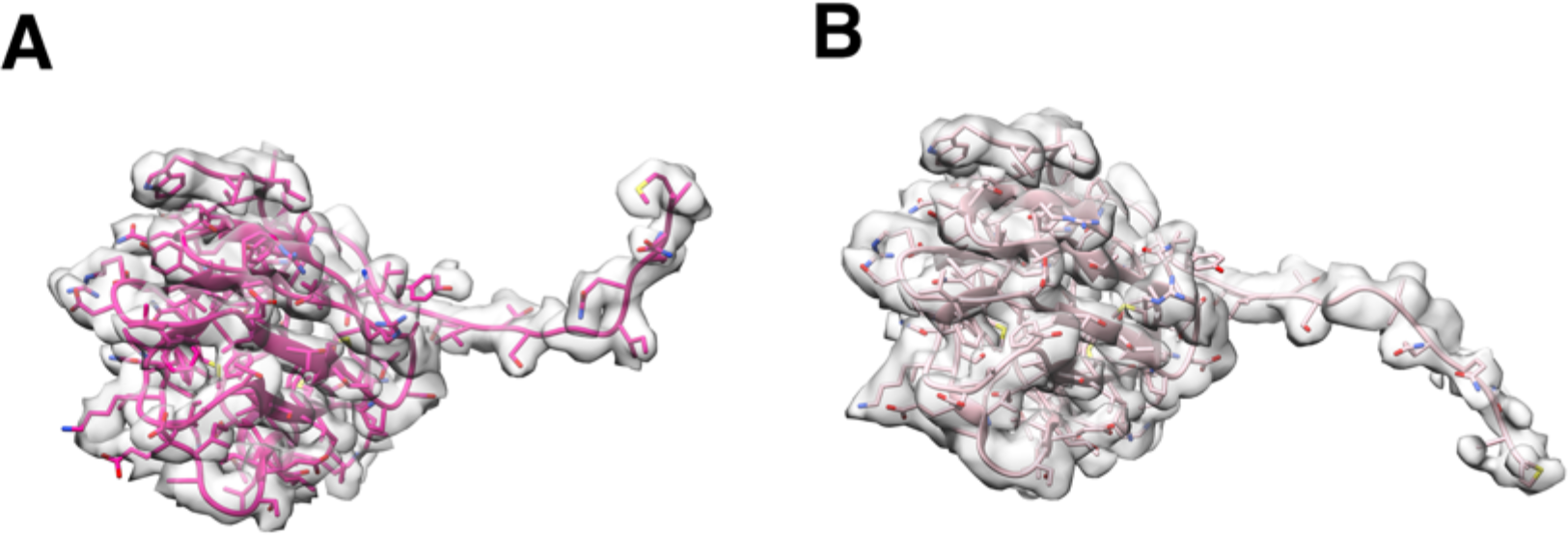
Two conformations of the P5 N-terminal base. (**A**) The typical conformation showing the N-terminal region that hug the neighboring subunit of the penton base. (**B**) Special conformation of the N-terminal region that can be wedged in-between the P3 trimers. The low occupancy of the special N-terminal conformation leads to poor map intensities due to map sharpening. So, the unsharpened filtered maps are used in the above image to fit the models.

**Figure 5–figure supplement 1.**
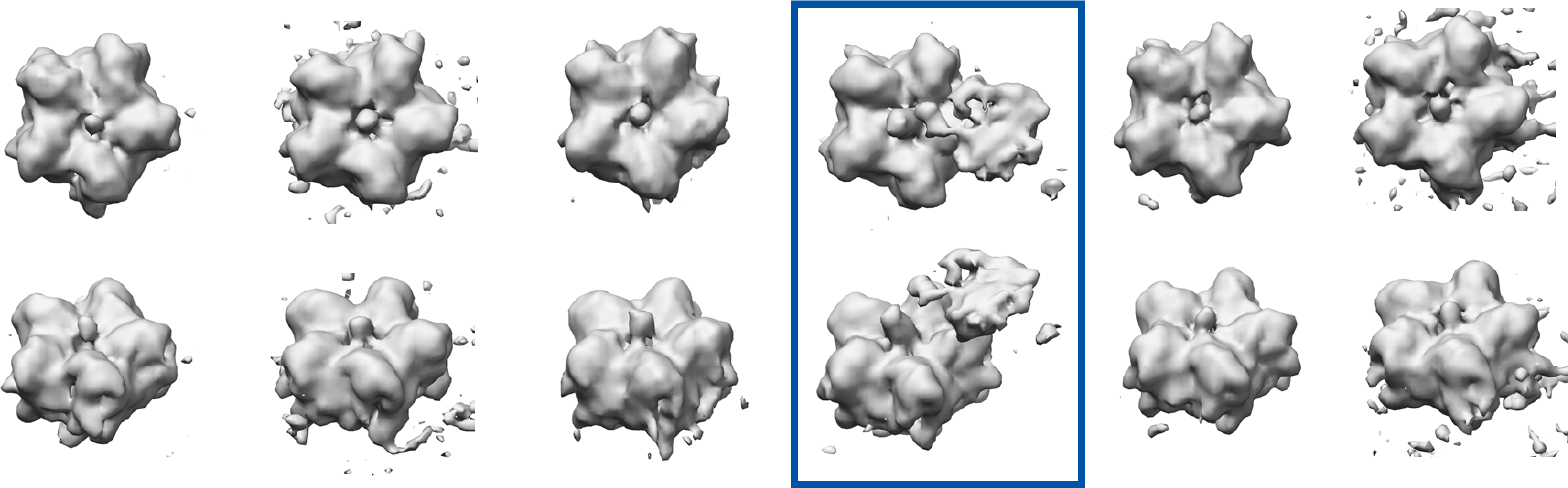
Top row shows all the maps that were generated by the localized reconstruction followed by initial 3D classification without image alignment and the bottom row shows the 45° tilted counterparts of the maps displayed in the top row. The map highlighted by the blue rectangle shows the class that was selected and the 3D classification was further refined with local image alignment. The resolution of the localized reconstruction was about 10Å. No tight masks were used during the 3D classification to avoid any masking bias. This results in a lower resolution map but the maps are more reliable.

**Figure 5–figure supplement 2.**
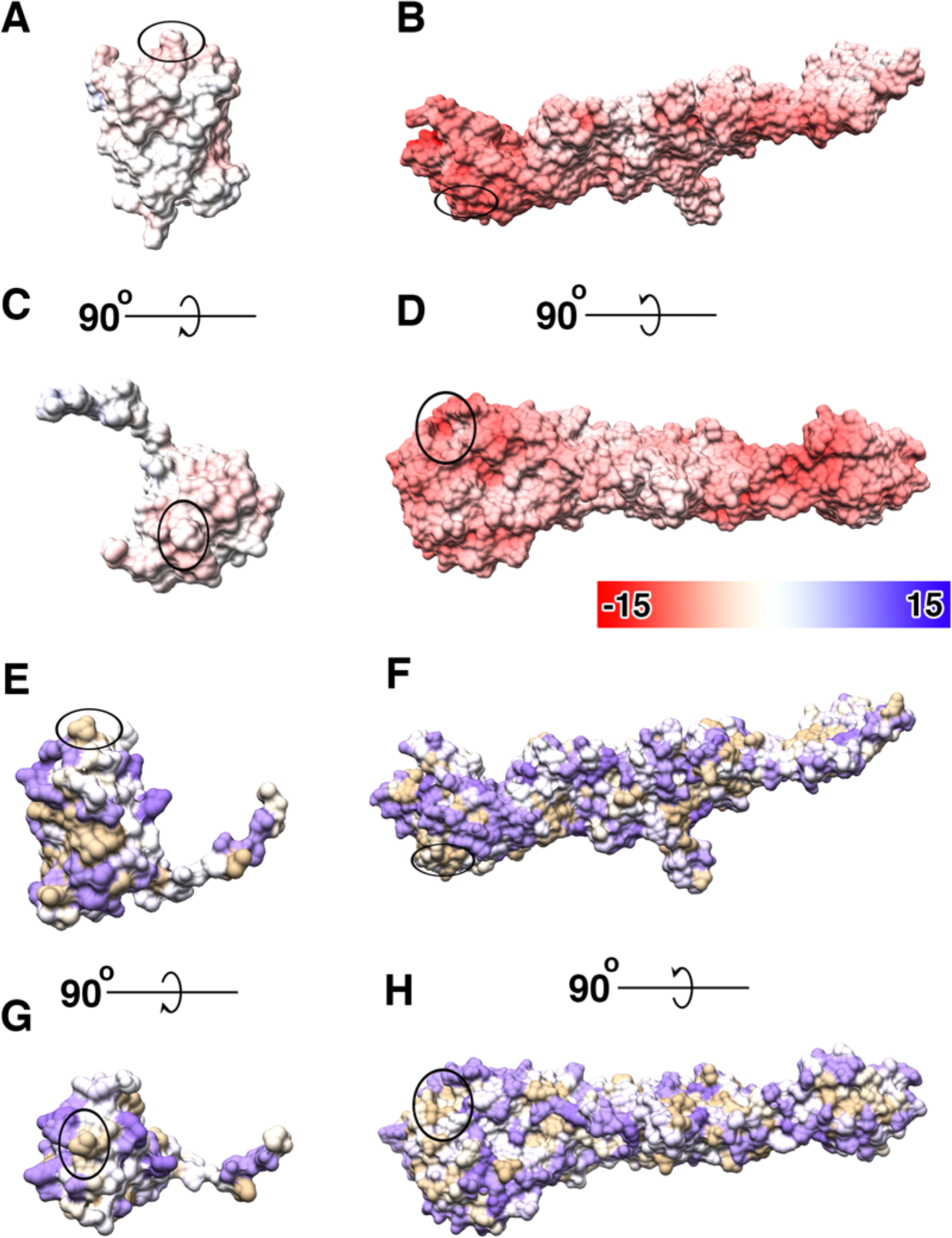
(**A**-**D**) Shows the distribution of coulombic electrostatic potential on the surface of P2 and P5 where red represents negative potential, blue represents positive potential and white for neutral. (**E**-**H**) Show the distribution of hydrophobic residues on the surface of P2 and P5 (according to Kyte and Doolittle scale) where purple represents least hydrophobic residues, brown represents most hydrophobic residues and white for neutral residues. The highlighted regions on all the images shows the region of interaction between the N-terminal domain of P5 and P2.

**Figure 5–movie supplement 1.**
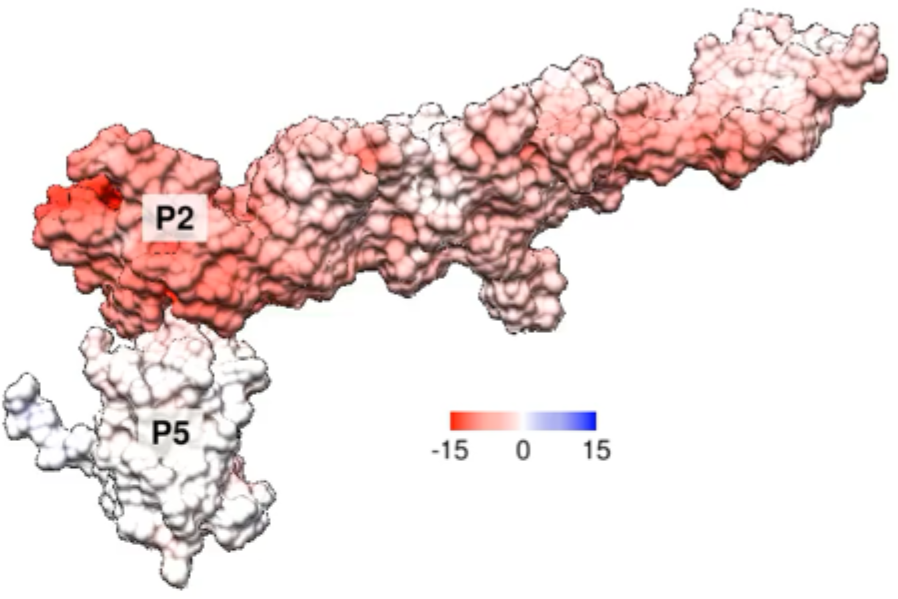
Shows the distribution of coulombic electrostatic potential on the surface of P2 and P5 and the regions of interaction between the N-terminal domain of P5 and P2.

**Figure 6–figure supplement 1.**
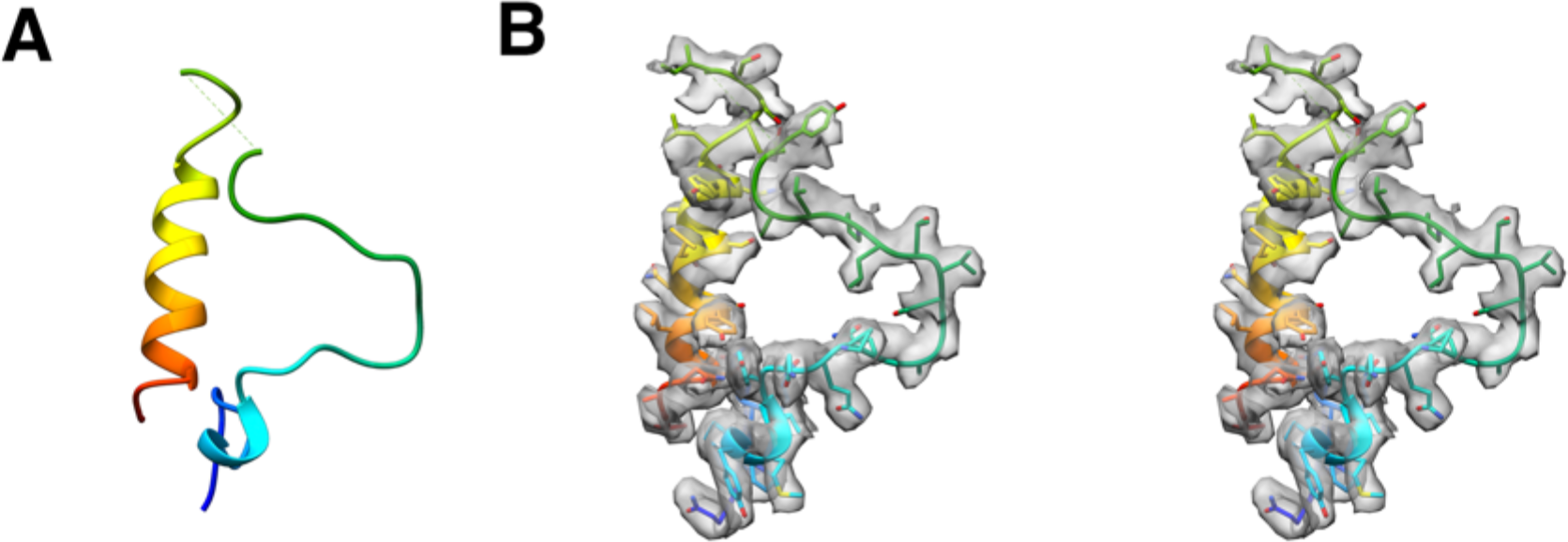
(**A**) The model of P16. (**B**) The stereo images of the model fitted into the CryoEM map.

**Figure 6–movie supplement 1.**
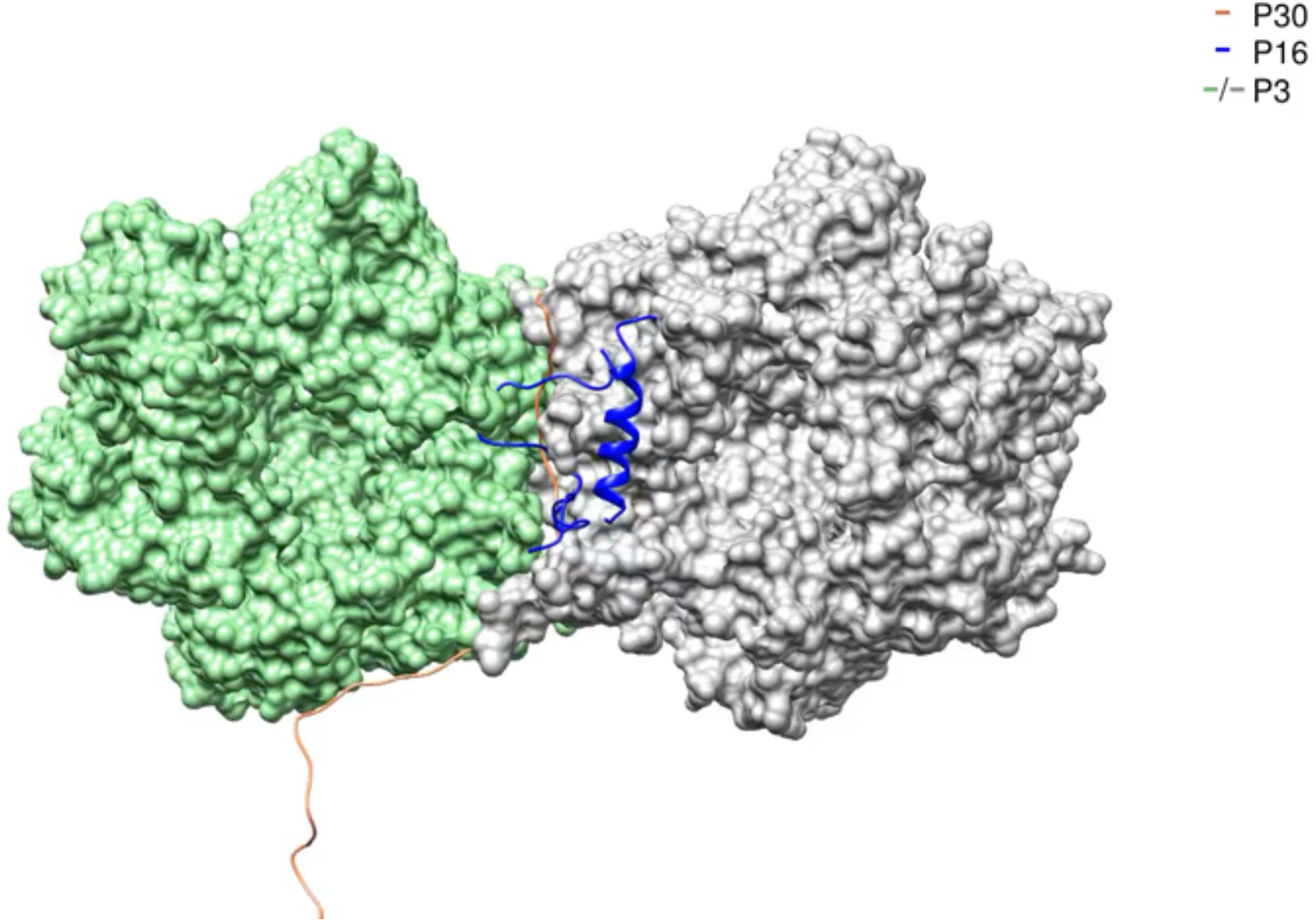
Movie showing the P3-P30-P16 complex.

## Supplementary Information: Figures

**Supplementary Figure 1.**
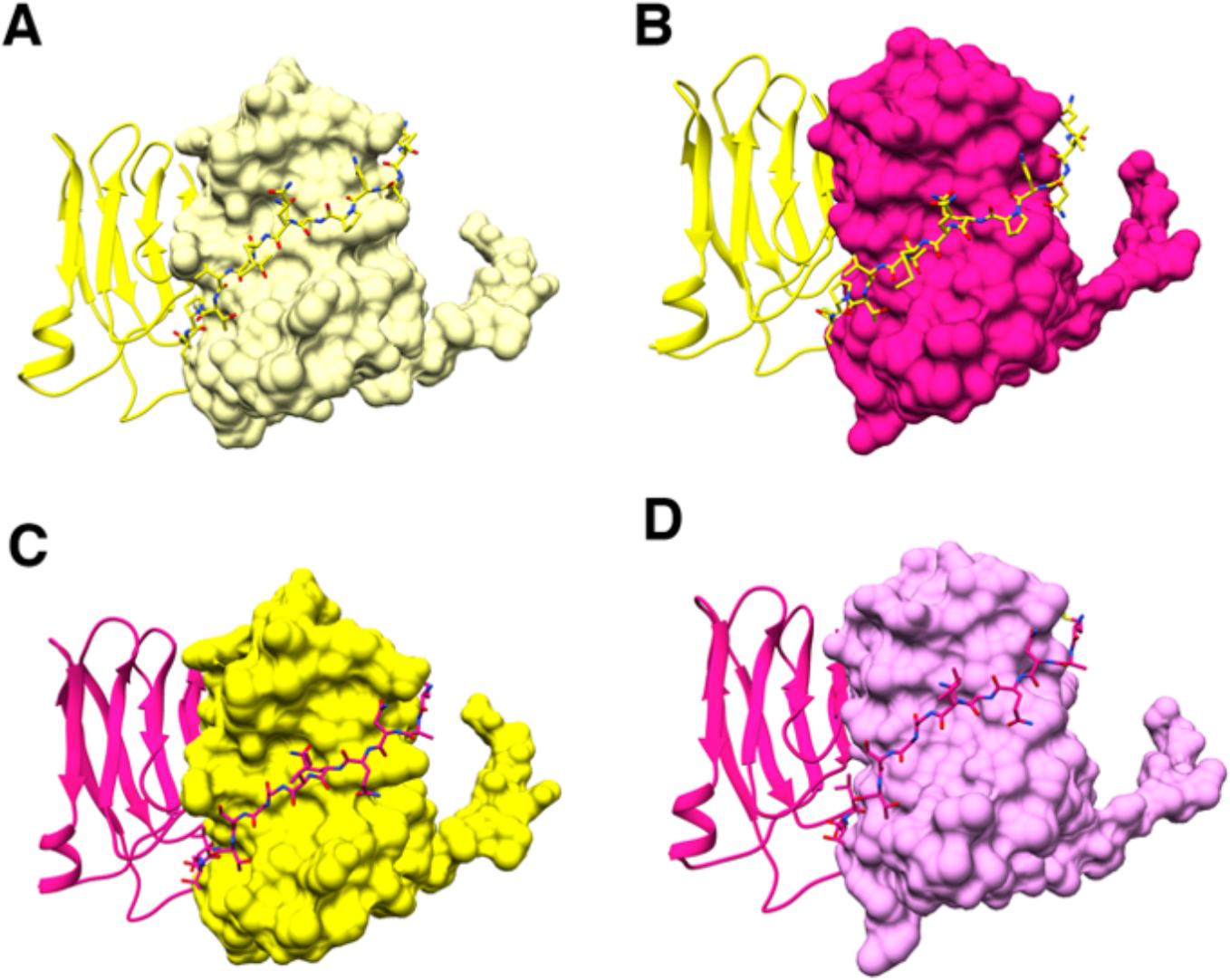
(**A**-**D**) Shows inter-digiting and hugging of the N-terminal residues of P31 (shown in shades of yellow) and P5 (shown in shades of pink) that stabilize the penton.

**Supplementary Figure 2.**
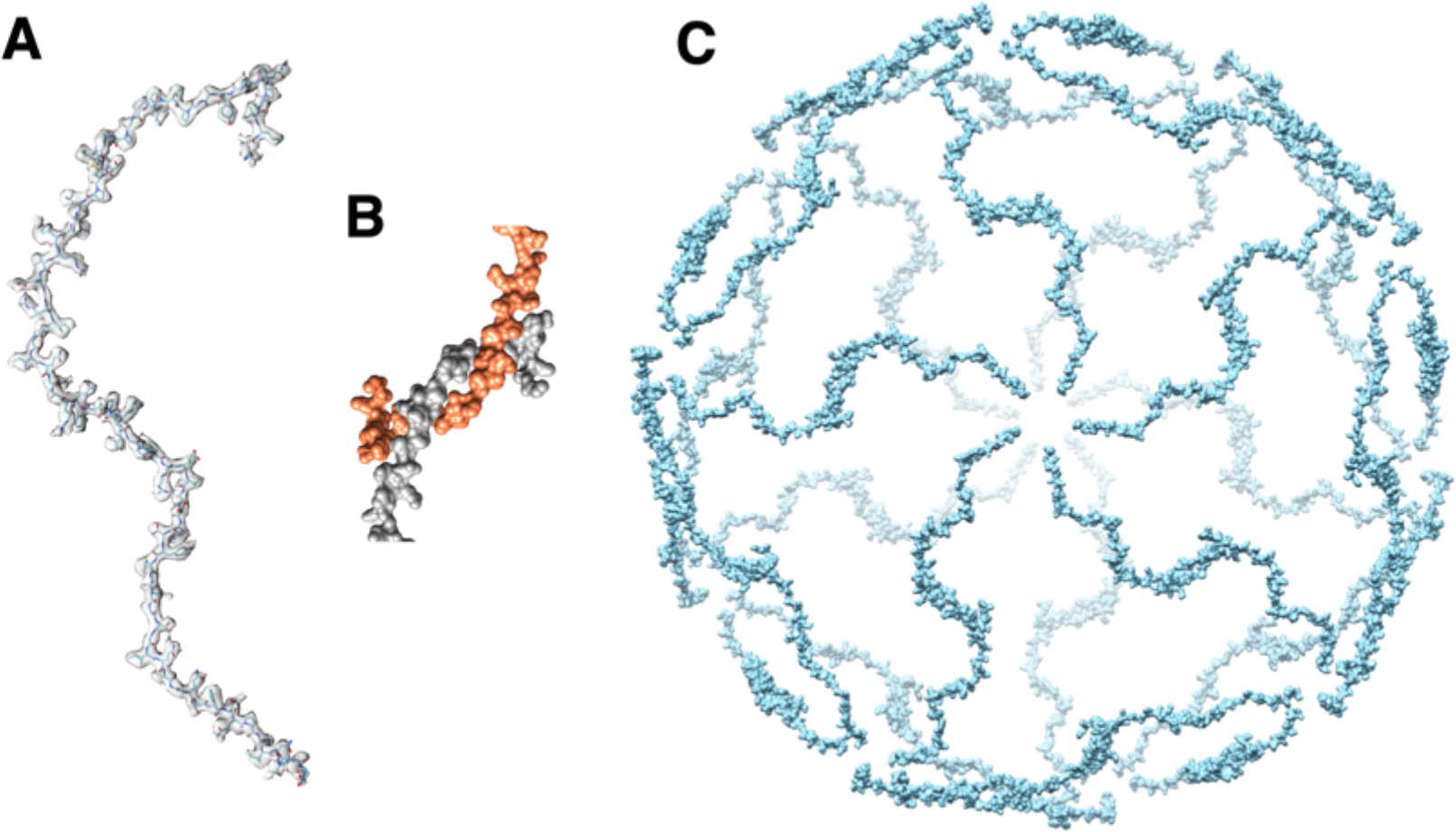
**A**) P30 monomer and its fit into the CryoEM density map. **B**) interlocked N-terminal hook forming the dimer that spans between 2 adjacent vertices. Residues Met1-Val32 are involved the formation of the N-terminal hook. **C**) Complete P30 cage that stabilizes the trisymmetrons and in turn the whole particle.

**Supplementary Figure 3.**
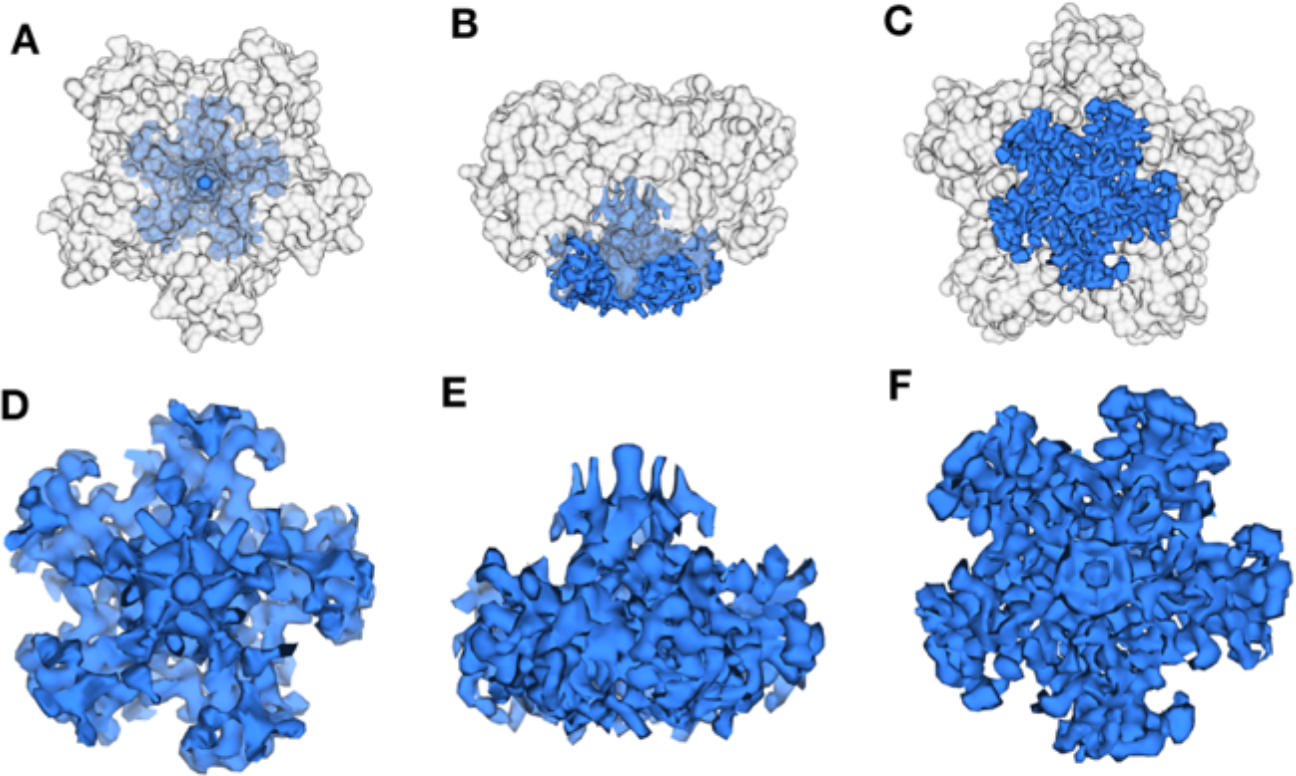
(**A**-**C**) Difference density map (blue) at the vertex region where the densities of the penton (P31 and P5) are shown in transparent grey. (**A, D)**,(**B, E**) and (**C, F)** show the top, side and the bottom view of the difference density map respectively.

**Supplementary Figure 4.**
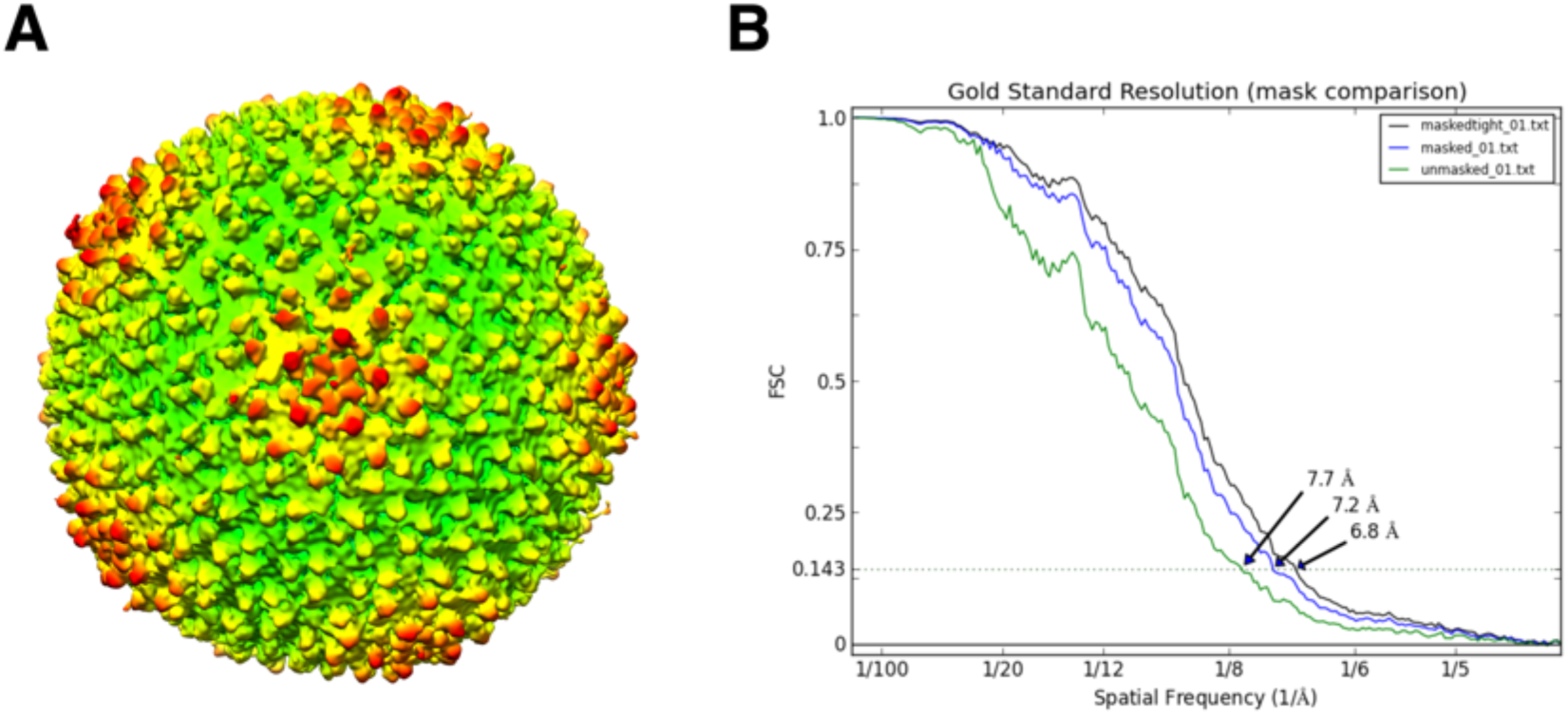
Asymmetric reconstruction of the wild type PR772 by symmetry breaking using EMAN2. (**A**) Preliminary map generated by the asymmetric reconstruction of PR772. Note that all the vertices show the heteropentameric penton. (**B**) FSC curve for the reconstruction.

**Supplementary Figure 5.**
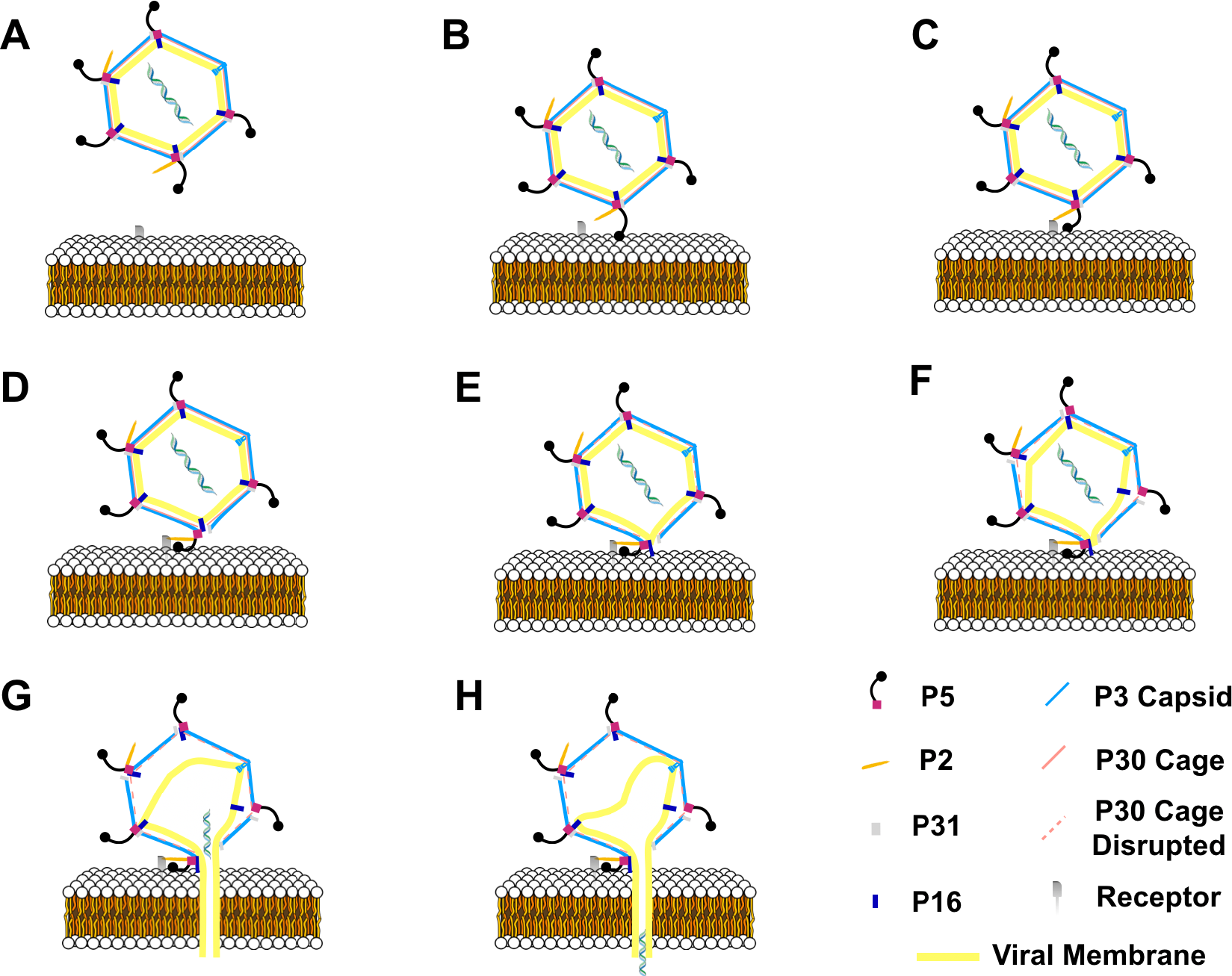
Schematic of the infection mechanism model of bacteriophage PR772. (**A**) Shows bacteriophage PR772 approaching the host membrane with the receptor. (B) The trimeric knob domain of P5 recognizes the host. The binding of the trimeric knob of P5 is transient. (**C**) High-affinity P2 binding to the surface receptor stabilizes the host binding. The binding is irreversible and attaches the virus to the host membrane. (**D**) P2 binding disturbs of the vertex complex by pulling the P5. (**E**) The disordered vertex complex disrupts the interactions between P5 and P30 resulting in a cascade that leads to the disruption of the P30 cage that stabilizes the viral capsid. (**F**) Disrupted P30 cage, destabilizes other adjacent vertices and the interactions that anchor the viral membrane. (**G**) P16 anchors into the host membrane and facilitates the formation of the membranous tube. (**H**) Destabilized viral membrane collapses and the dsDNA is delivered into the host.

**Supplementary Figure 6.**
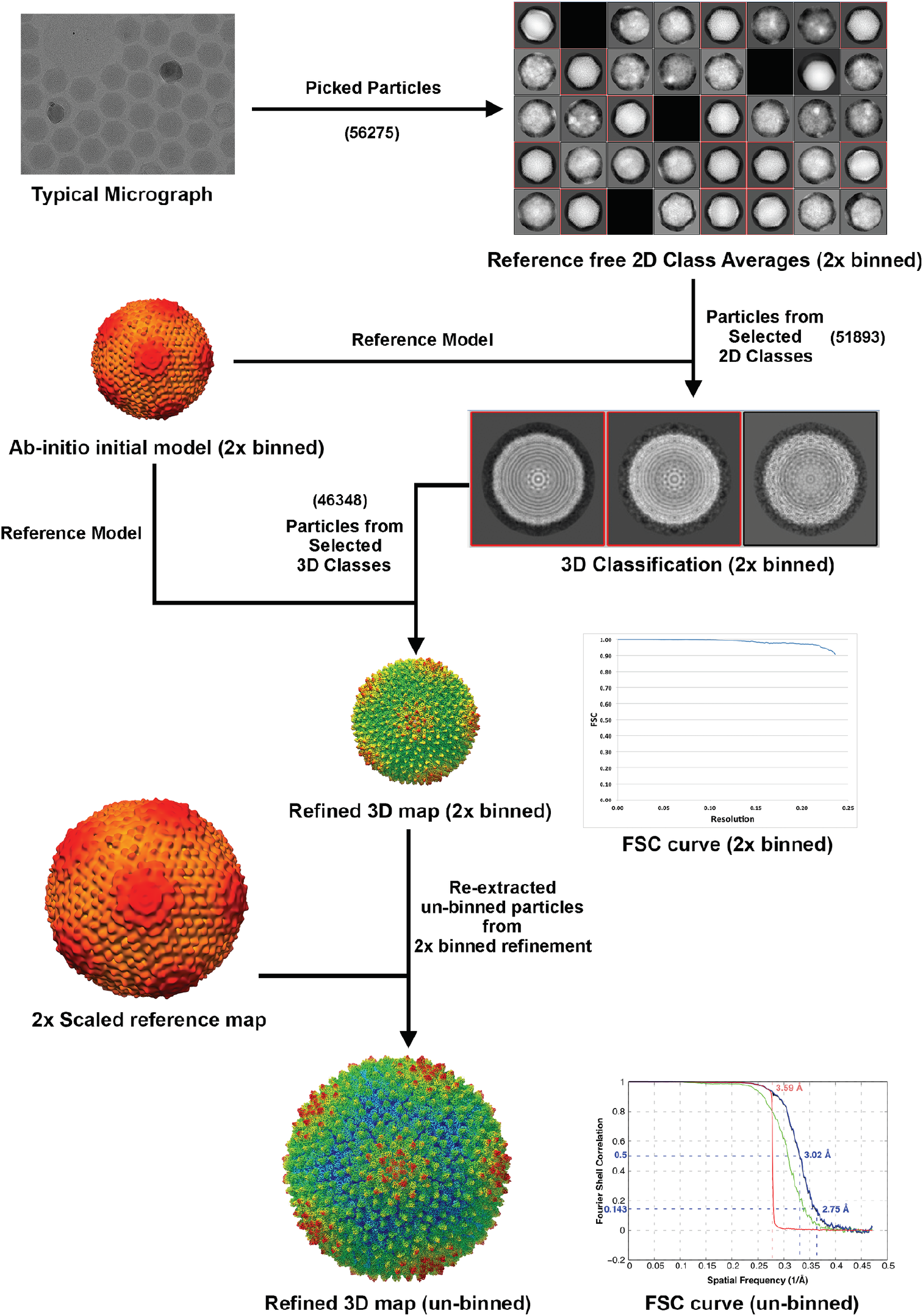
Flowchart of the 3D reconstruction of the icosehadrally averaged PR772 map. The preprocessed micrographs were used to autopick the particles using template matching. The autopicked particles were extracted by 2x bining. The good particles were selected by inspecting the 2D class averages from the reference free 2D classification. An icosahedrally averaged ab-initio initial model was generated and used as a reference to do 3D classification. The particles from the dominant classes were selected and 3D refined. On reaching Nyquist, the particles from the 2x binned reconstruction were re-extracted without binning for further 3D refinement. The reference map was scaled accordingly.

## Supplementary Information: Text

### Supplementary Text 1

Comparison of RMSD of different protein subunits from PR772 and PRD1, from a similar region of the map using Chimera and SuperPose. Sequence-guided structure alignment of most proteins from PR772 and PRD1 yielded poor RMSD values even though they have high sequence identity. So, secondary-structure based alignment (without the use of sequence alignment) was also performed. We see a significant variation in the models for P31, P30, P16 and minor variations in overall P3 structure. The P31 protein from PR772 has 100% protein sequence identity with P31 from PRD1, but the model comparison reveals a RMSD of 16.7Å with significant registry error in the model from PRD1. P30 has a 97.6% sequence identity and the model comparation reveals a RMSD of 4 Å. P16 with a sequence identity of 96%, has a RMSD of 5.49 Å. P3 has a 99.7% protein sequence identity and the model comparation shows an overall RMSD of 1Å but the most crucial and the functionally important C-terminal and N-terminal regions show a RMSD of 2.3Å and 3.0Å respectively.

Below are the protein alignments.

### P31

Protein Sequence Identity: 100%

### Sequence-Guided Structure Alignment

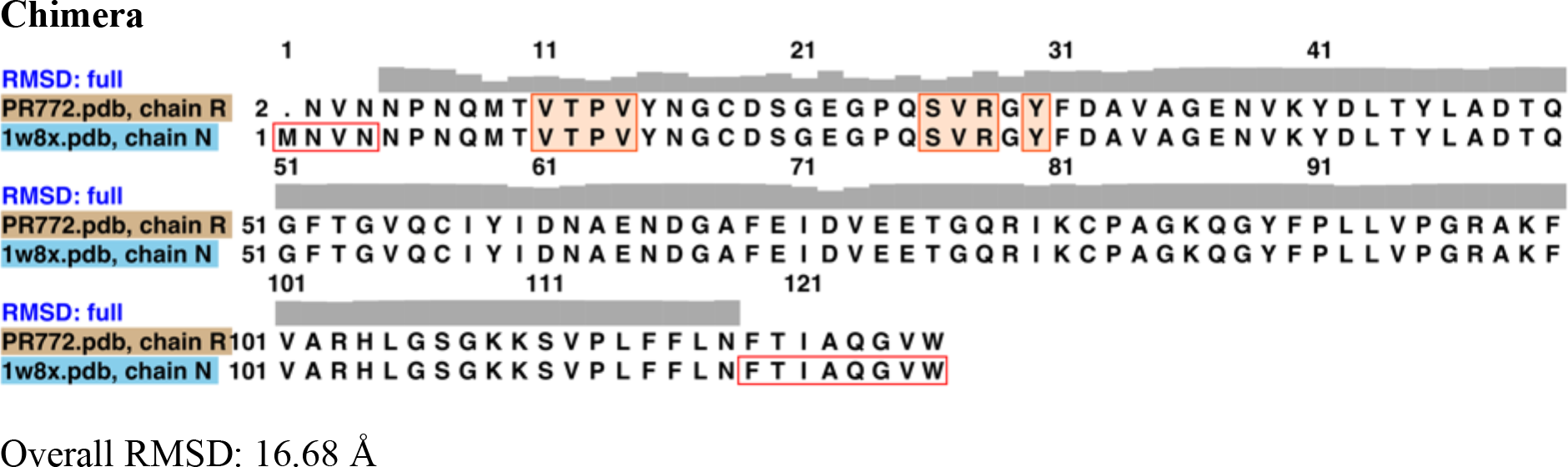

### SuperPose

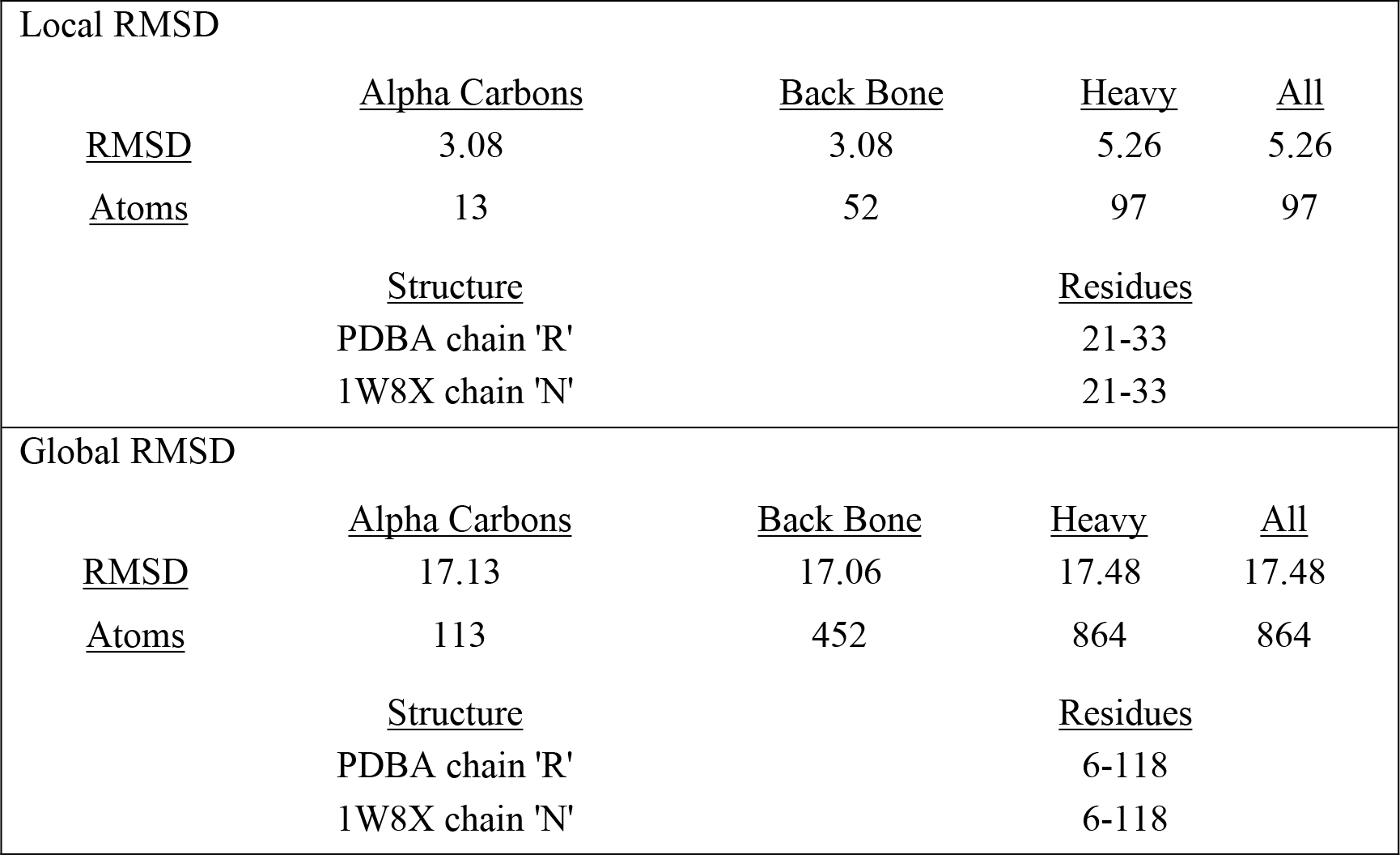

### Secondary Structure Based Alignment

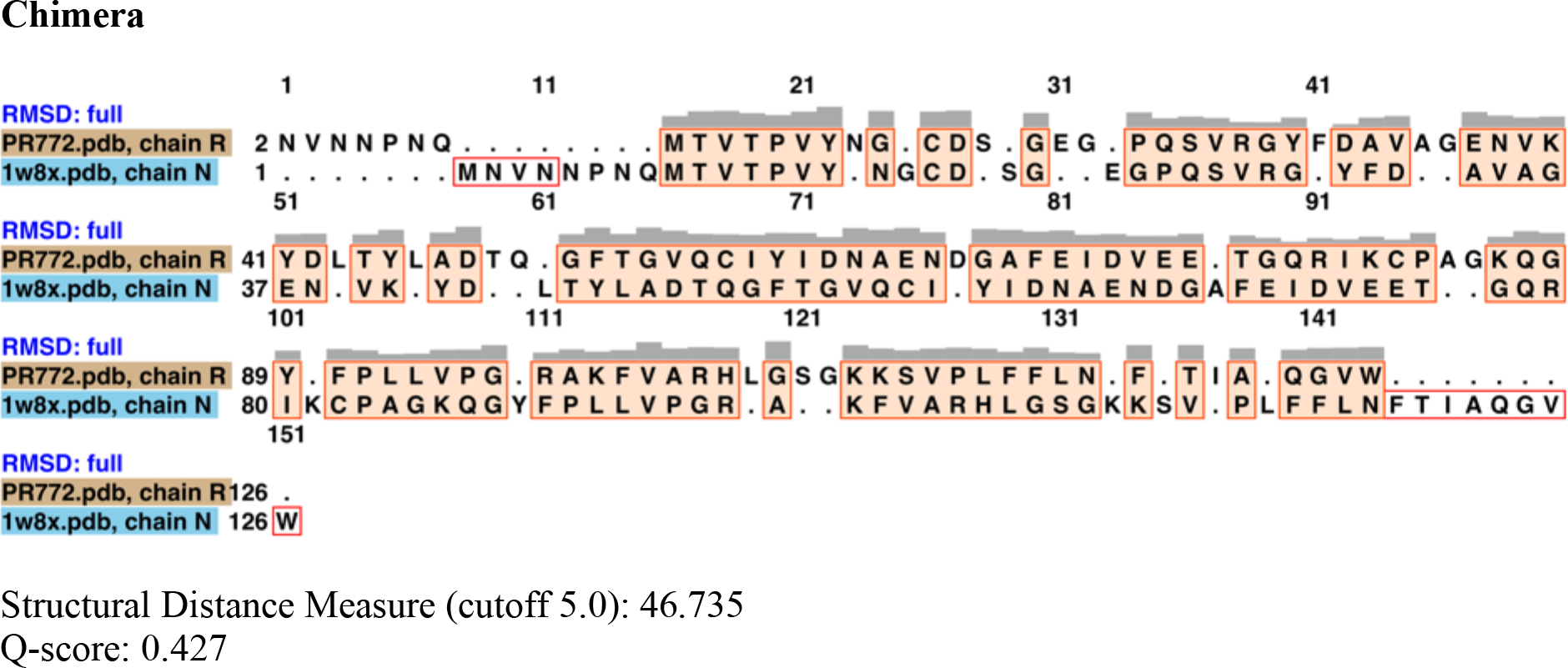

### P30

Protein Sequence Identity: 97.6%

### Sequence-Guided Structure Alignment

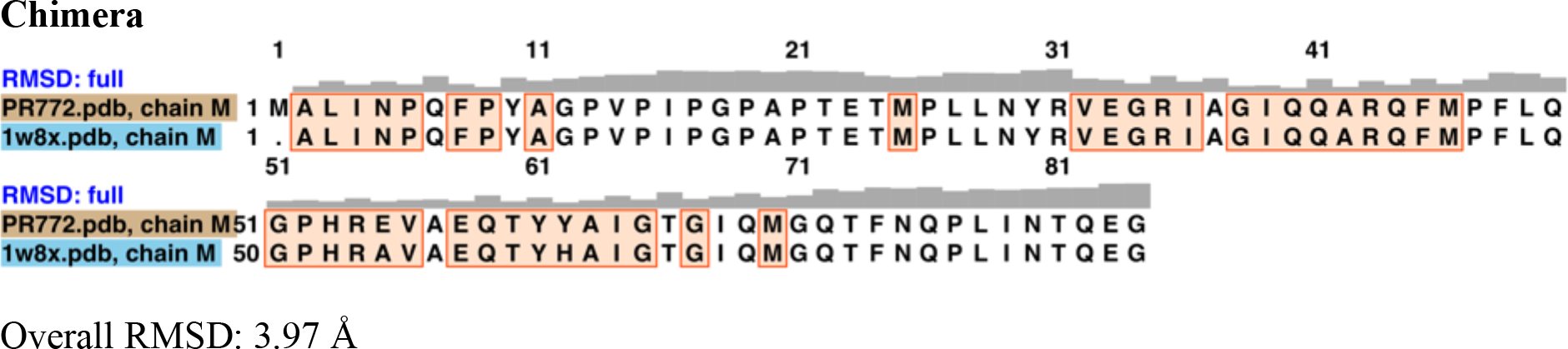

### SuperPose

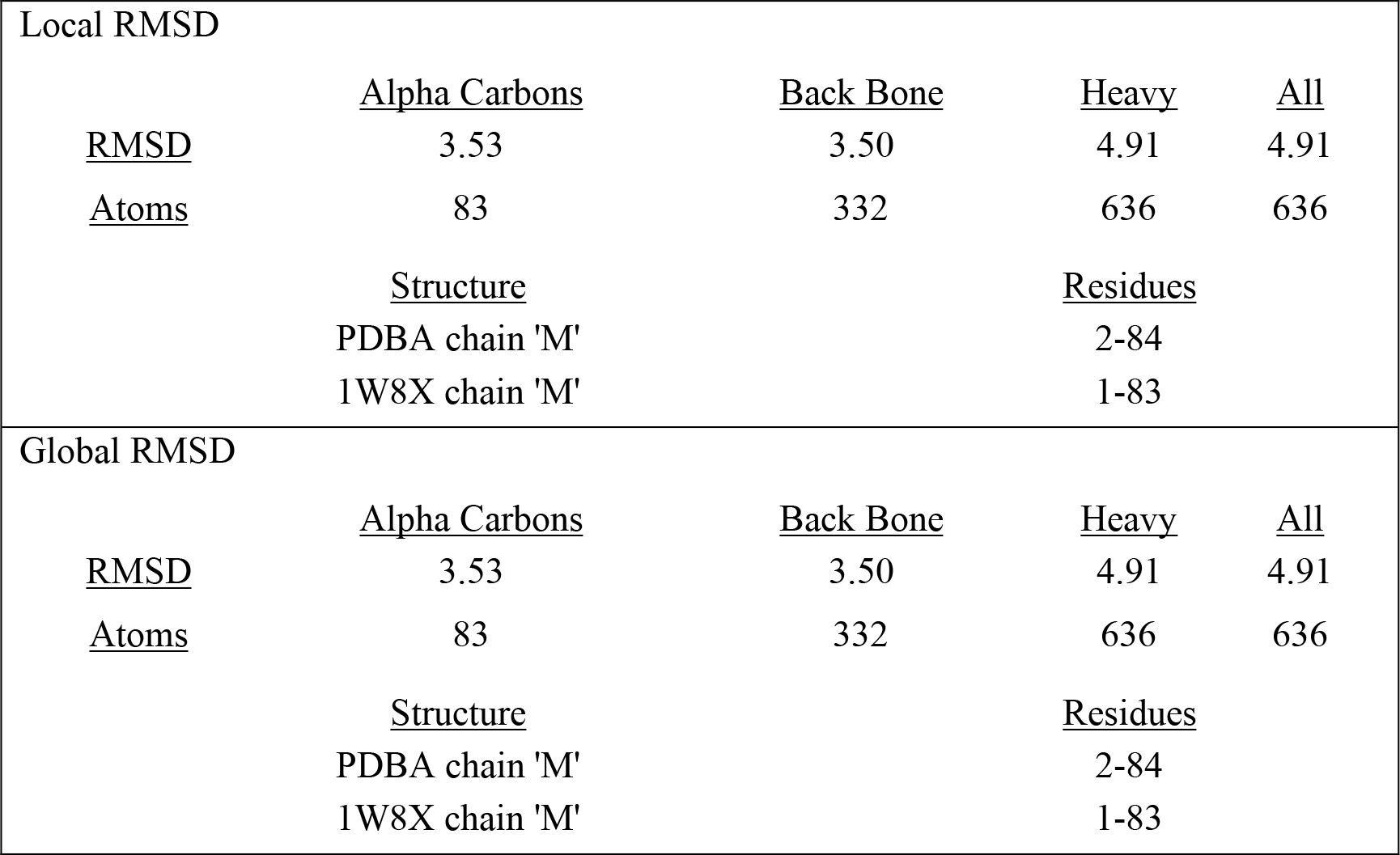

### Secondary Structure Based Alignment

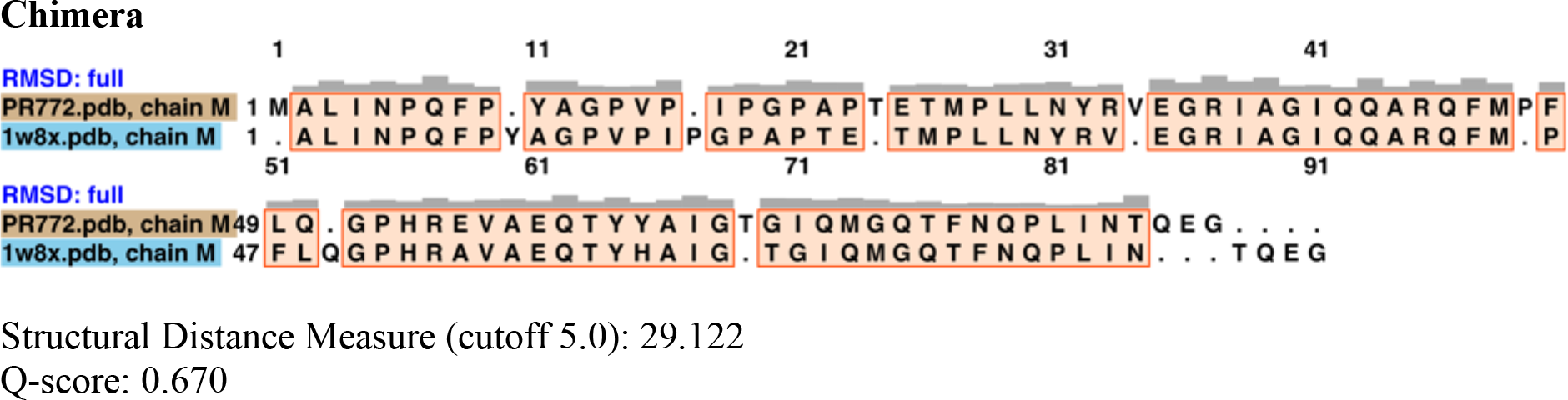

### P16

Protein Sequence Identity: 94%

### Sequence-Guided Structure Alignment

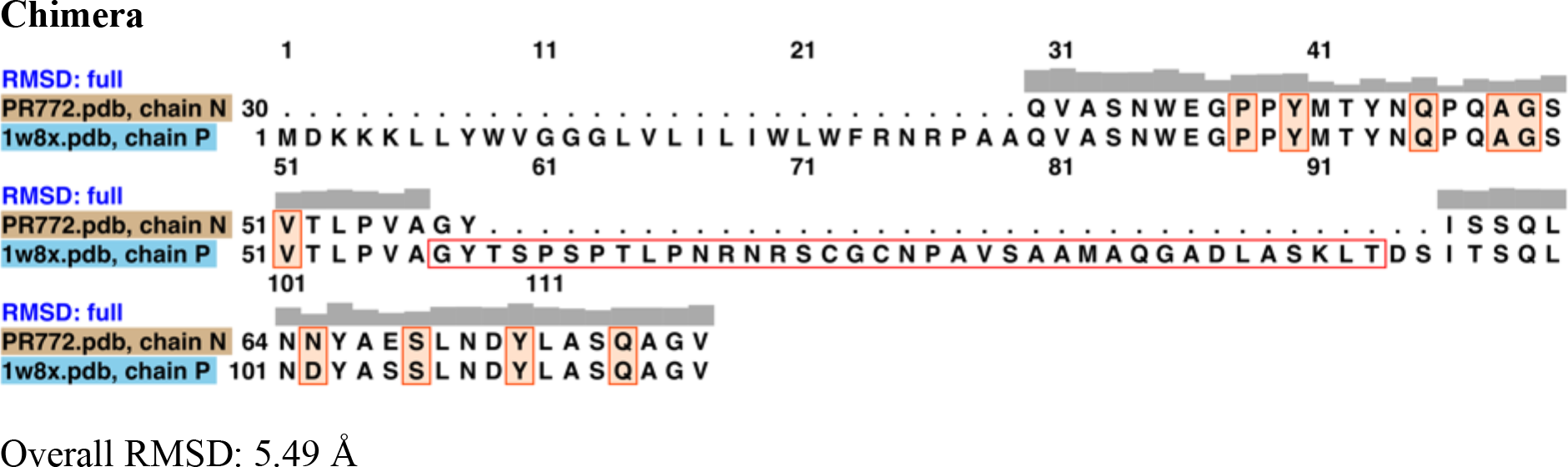

### SuperPose

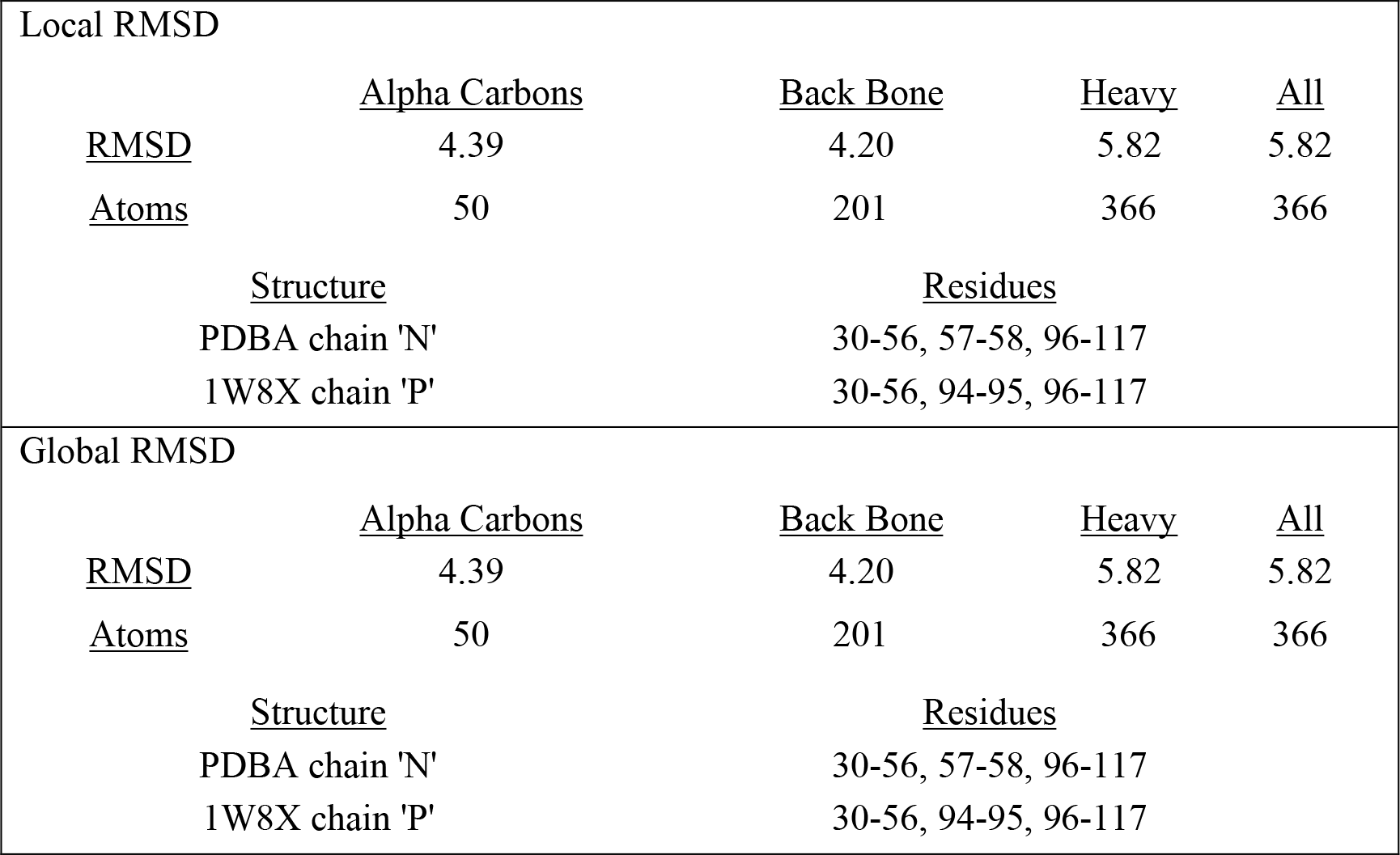

### Secondary Structure Based Alignment

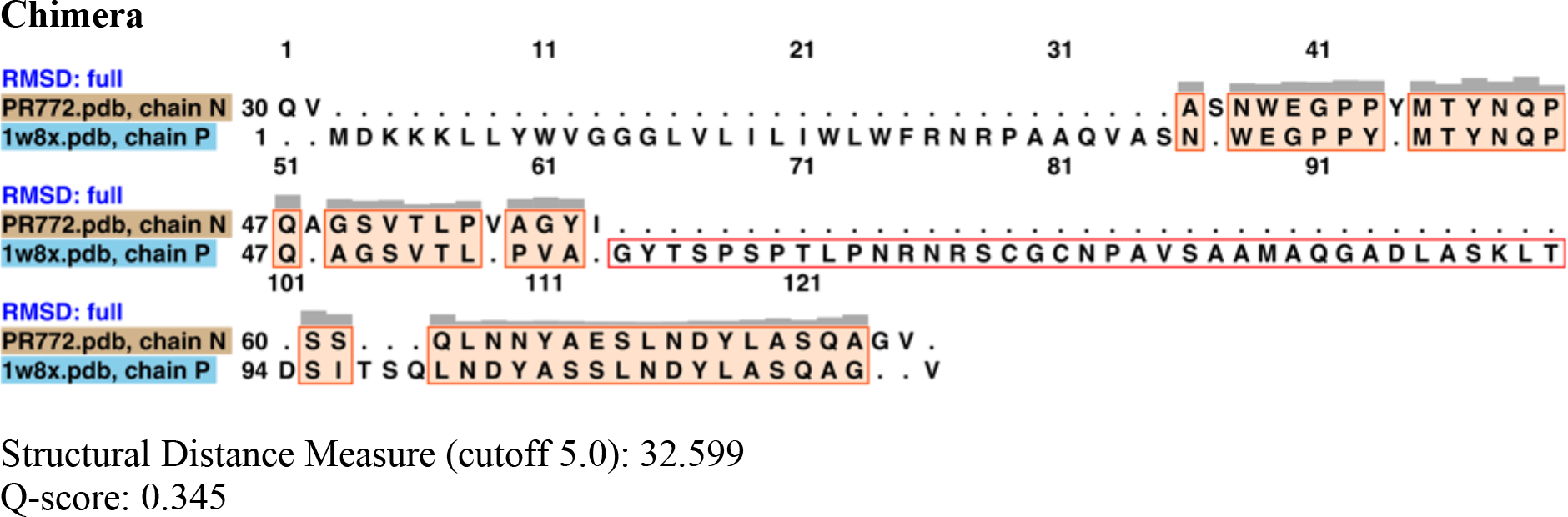

### P3 (Typical Case)

Protein Sequence Identity: 99.7%

### Sequence-Guided Structure Alignment

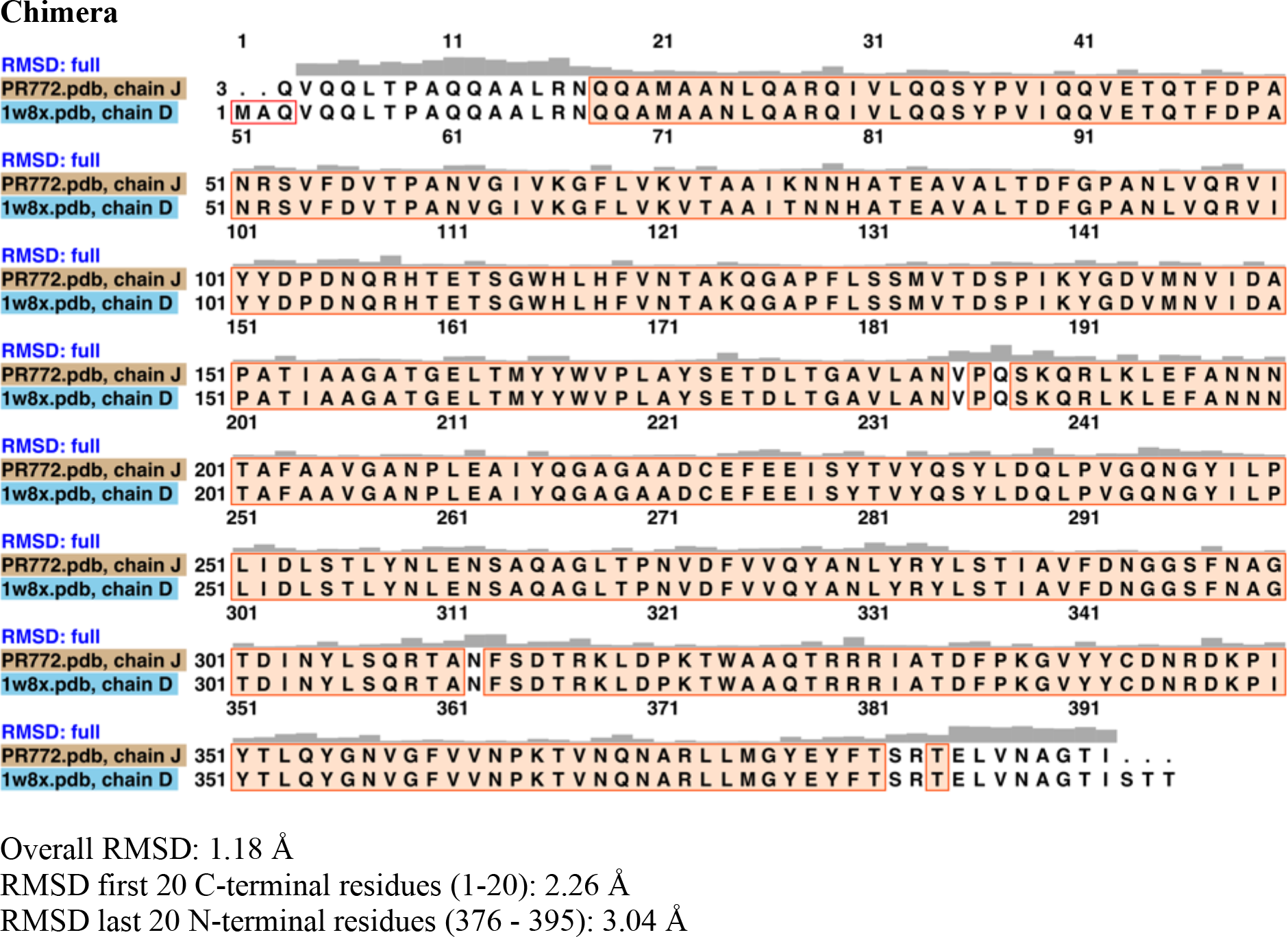

### SuperPose

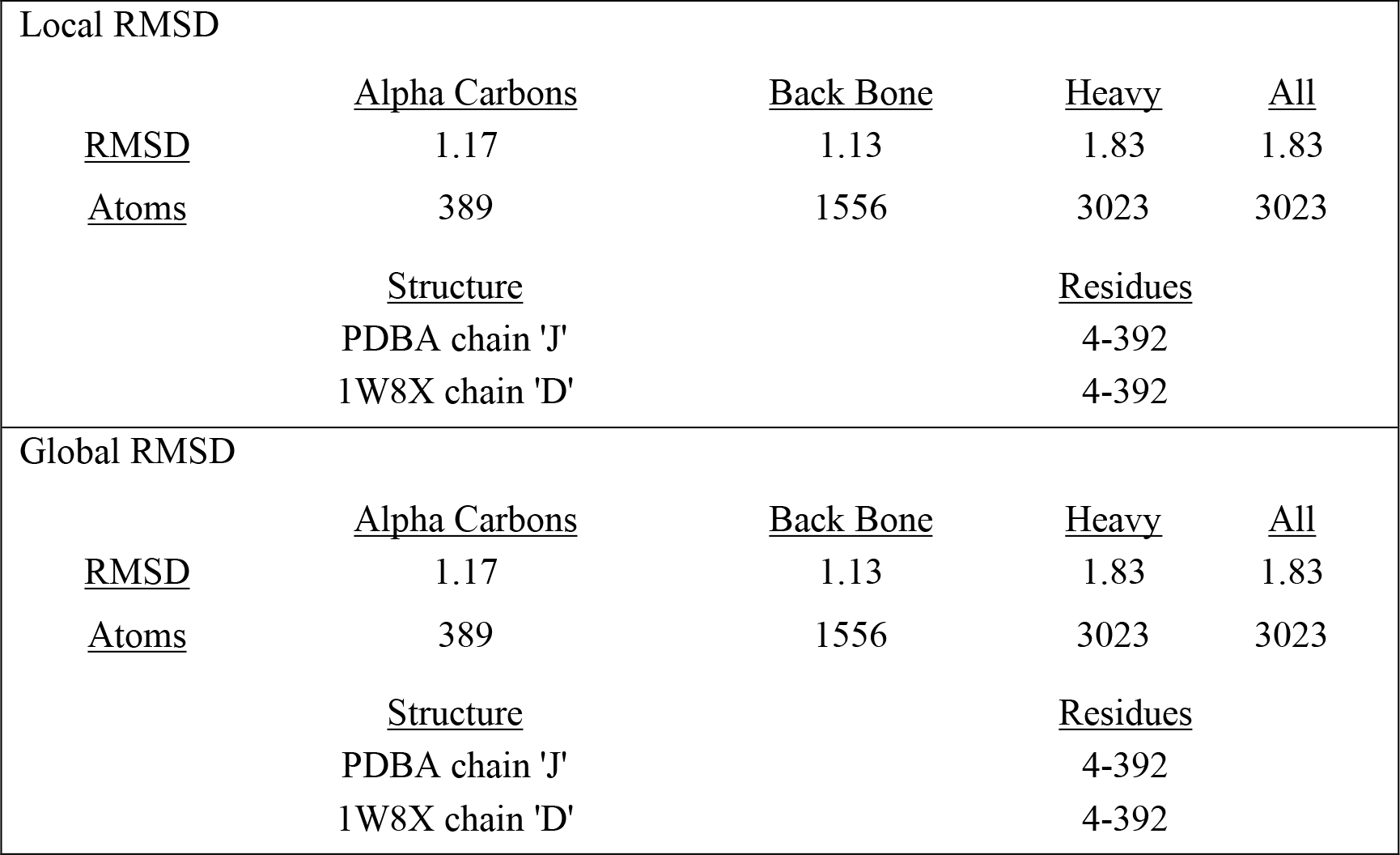

SuperPose difference distance map for P3

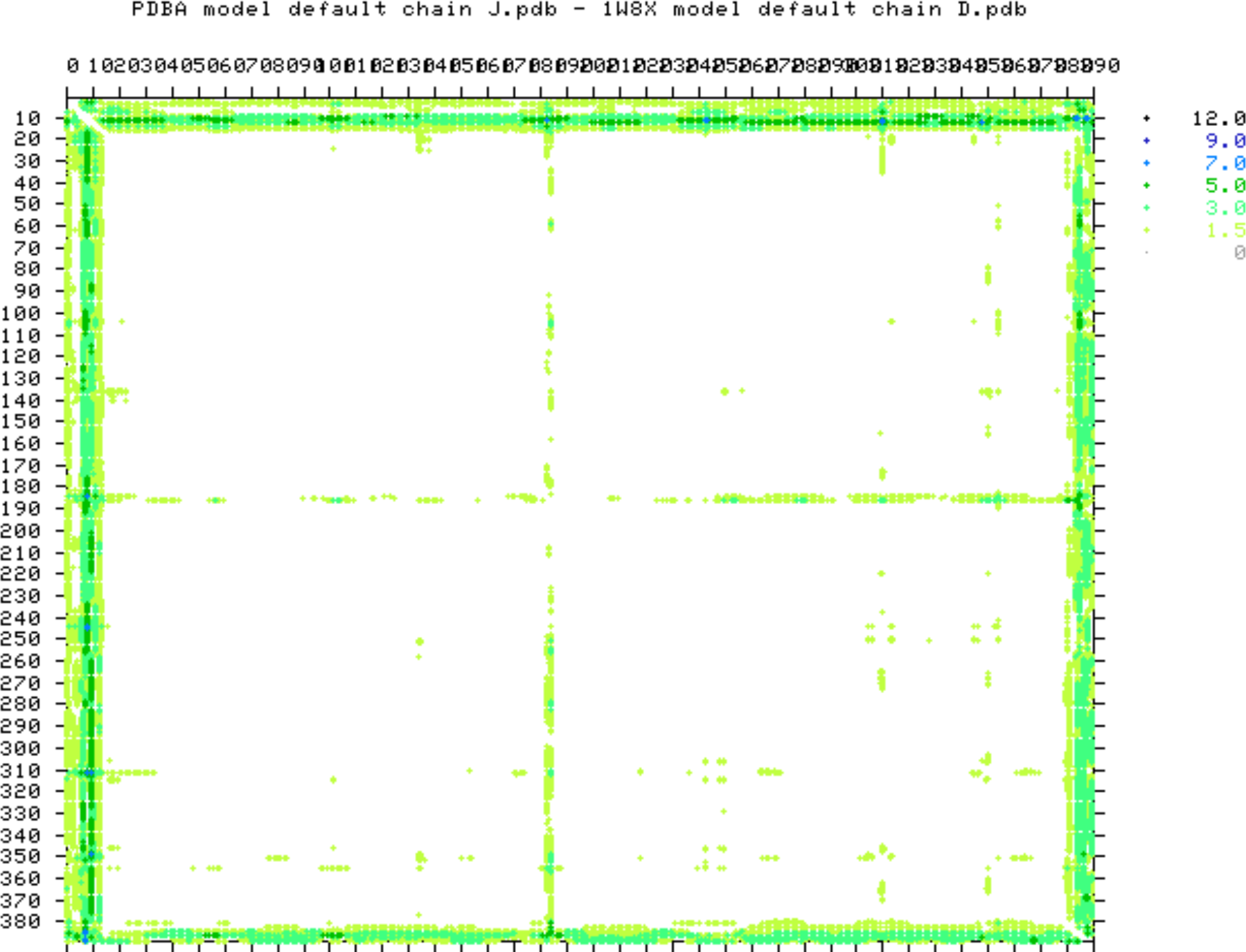

### Secondary Structure Based Alignment Chimera

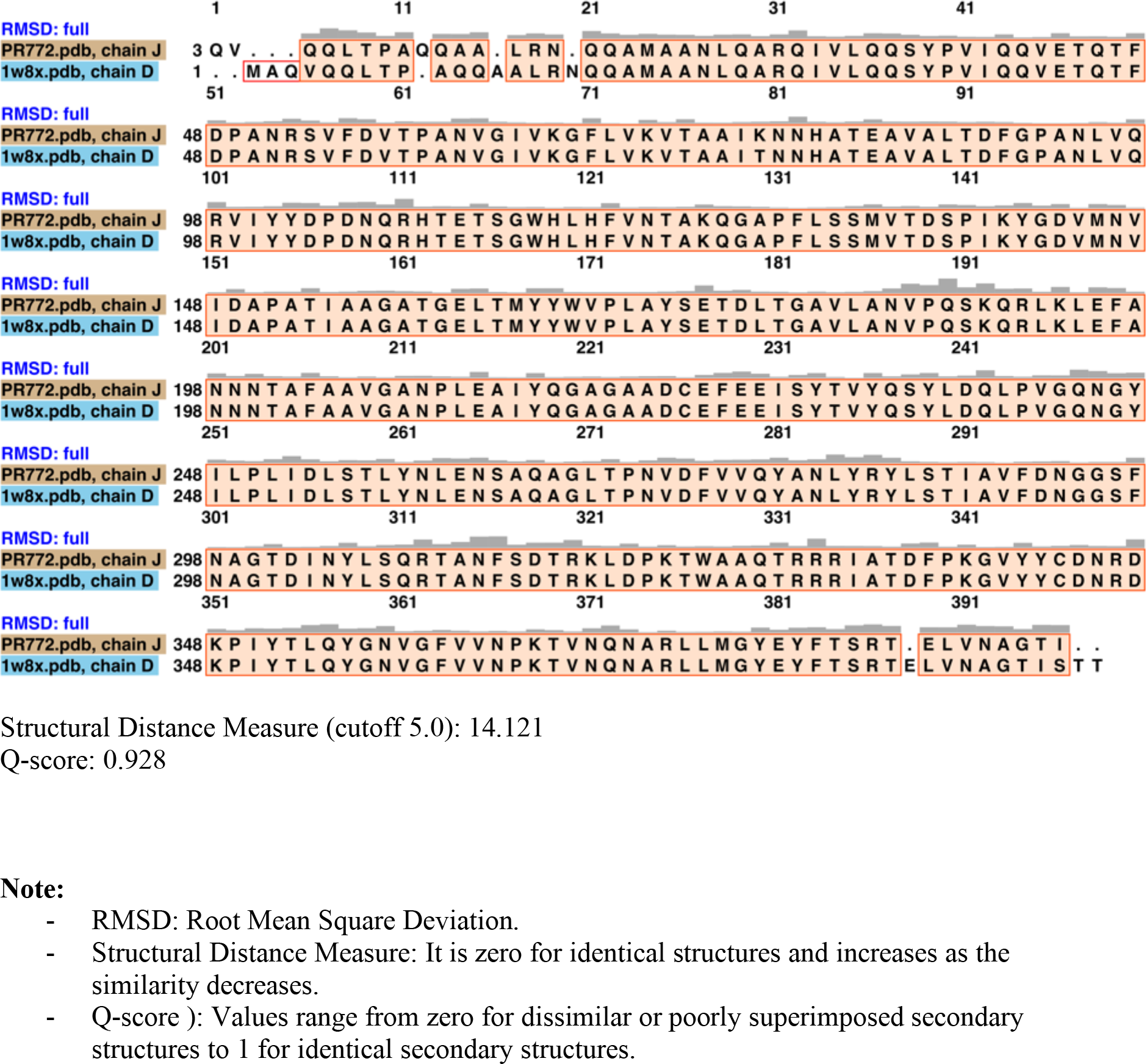

### Supplementary Text 2. Mtriage summary of the map quality analysis

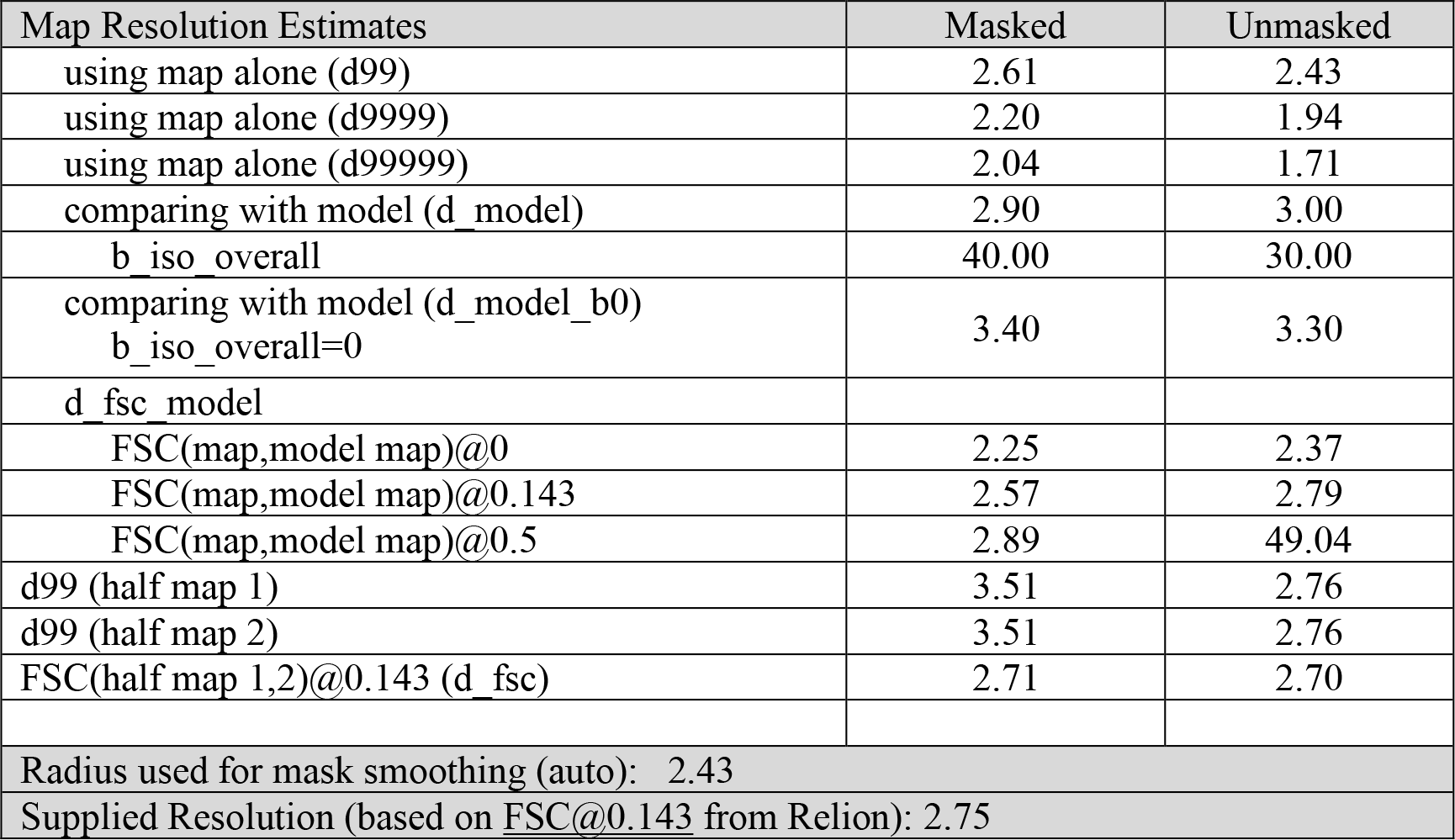

### Map-model CC (overall)

|  |  |
| --- | --- |
| CC_mask | 0.8349 |
| CC_volume | 0.8348 |
| CC_main_chain | 0.8386 |
| CC_side_chain | 0.8085 |

Legend:

- **d_fsc** - highest resolution at which the experimental data are confident. Obtained from FSC curve calculated using two half-maps and taken at FSC=0.143.
- **d99** - resolution cutoff beyond which Fourier map coefficients are negligeably small. Calculated from the map.
- **d_model** - resolution cutoff at which the model map is the most similar to the target (experimental) map.
- **d_FSC_model** - resolution cutoff up to which the model and map Fourier coefficients are similar.

